# Gene Birth in a Model of Non-genic Adaptation

**DOI:** 10.1101/2022.07.31.502179

**Authors:** Somya Mani, Tsvi Tlusty

## Abstract

**Background:** Over evolutionary timescales, genomic loci can switch between functional and non-functional states through processes such as pseudogenization and *de novo* gene birth. Particularly, *de novo* gene birth is a widespread process, and many examples continue to be discovered across diverse evolutionary lineages. However, the general mechanisms that lead to functionalization are poorly understood, and estimated rates of *de novo* gene birth remain contentious. Here, we address this problem within a model that takes into account mutations and structural variation, allowing us to estimate the likelihood of emergence of new functions at non-functional loci.

**Results:** Assuming biologically reasonable mutation rates and mutational effects, we find that functionalization of non-genic loci requires the realization of strict conditions. This is in line with the observation that most *de novo* genes are localized to the vicinity of established genes. Our model also provides an explanation for the empirical observation that emerging proto-genes are often lost despite showing signs of adaptation.

**Conclusions:** Our work elucidates the properties of non-genic loci that make them fertile for adaptation, and our results offer mechanistic insights into the process of *de novo* gene birth.

## Background

An organism’s genome consists of several types of loci that are known to perform diverse functions: these functional loci include genes, gene regulatory regions, and sequences maintaining chromosome structure [1]. However, measuring the proportions of the functional parts of genomes is a subject that is fraught with disagreements, which stem from problems with our conception of what constitutes a functional genomic locus [2]. While evidence of selection at a genomic locus is a strong indicator of function, problems arise when one considers the emergence of functionality in a previously non-functional locus. The subject of emergence of functionality is especially relevant for the study of *de novo* gene birth, whereby novel genes are born from previously non-genic loci [3]. Specifically, *de novo* gene birth involves a previously non-genic locus gaining the ability to express a functional product. Hereafter, our definition of functional product includes both proteins and RNA.

*De novo* gene birth is a widespread phenomenon: well known examples of *de novo* genes exist across multiple evolutionary lineages, such as plants [4], fungi [5] and animals [6], including humans [7]. At the same time, measuring the rate of *de novo* gene birth is not straightforward [3]: Firstly, evolutionary conservation forms a basic criterion for a sequence to even be considered a ‘gene’, and candidates for *de novo* emerged genes necessarily show some level of conservation. Secondly, the only conclusive criteria for proving *de novo* emergence is to identify the non-coding ancestral sequence, and identify *enabler* mutations that first allowed expression of this sequence [8]. These two criteria are at odds with each other, since the likelihood that traces of the homologous ancestral sequence are lost increases with time, and hence, with conservation level. Therefore, strict criteria for choosing candidate ‘genes’ and strict criteria to ascertain ‘*de novo* emergence’ lead to an undercount [9]. On the other hand, genomic analyses based on synteny [10] and phylostratigraphy [11] find indirect evidence of *de novo* gene emergence based on the lack of a homologous ancestral coding sequence. These indirect counts are most likely to be overcounts, and thus the proposed prevalence of *de novo* gene birth remains contested [12]. In parallel, new methods to detect *de novo* genes based on machine learning techniques are also being developed [13].

Our understanding of the range of possible mechanisms of *de novo* gene birth comes from several well-known case studies. The proposed mechanisms are typically expressed as a series of sub-steps involving gains and losses of stable transcription, open reading frames [3], and sequences for nuclear export of RNA [14]. Case studies have also uncovered how certain genomic regions, such as introns [15], and certain tissues like the brains and testes of animals [3] favor the emergence of *de novo* genes. These mechanisms serve as very useful descriptors of the conditions under which *de novo* gene birth becomes likely. Nevertheless, in order to formulate a predictive theory of *de novo* gene birth, it is necessary to connect these complex descriptions with empirically measured rates and effects of simple genomic changes such as mutations and structural variations. In this work, we attempt to bridge this gap: We envisage that contributions of genomic loci to organismal fitness, which is often measured in terms of the organism’s growth rate, can be used as a correlate of the functionality of these loci. For example, in [16], the physiological activity of putative yeast proto-genes was demonstrated when overexpression of these genes impacted growth rate. In this study, we use an evolutionary model to explore how the contribution of a non-functional genomic locus to organismal fitness might increase over time due to accumulating mutations, and investigate the implications of this process for *de novo* gene birth.

The effects of novel, spontaneous mutations on growth rate have been experimentally measured for various organisms [17]. The outcomes of these studies are represented as DFEs (distributions of fitness effect), which are distributions of the magnitude of mutational effects on growth rate. In our model, we sample biologically reasonable distributions of fitness effect of mutations using recently measured DFE parameters for *Chlamydomonas reinhardtii* [18] as reference. Notably, observations in [18] indicate that the DFE of specific regions of the *C reinhardtii* genome, such as exons, introns or intergenic sequences, are similar to each other and to the DFE of the whole genome. However, in general, the DFE is known to vary across different regions of the genome [19], and across different species [20]. Particularly, we expect most mutations incident on non-functional genomic loci to be neutral. Accordingly, we sample a wide range of DFEs in the vicinity of the DFE reported in [18], and a large number of the sampled DFEs mostly produce neutral mutations.

Over a time scale of millions of years, in addition to small mutations (< 50bp), one can also expect large structural variations (from 50 bp up to several megabases) to impact the evolution of genomic loci [21]. While the rate of structural variation is estimated to be hundreds of times slower than the rate of small mutations, its effect is likely to be much larger [22]. Of particular importance to our question is the possibility that the entire genomic locus under consideration gets deleted. Therefore, we test in our model whether an initially non-functional locus can survive in the face of locus deletion for long enough to allow for adaptation.

Finally, we consider the particular case of *de novo* gene birth, whereby a previously non-functional locus gains the ability to express a functional product (protein/RNA). Recent studies report how new genes gain expression [23] and functionality [24] over time. Measurements from mutational scans of protein-encoding genes indicate that the overall fitness contribution of a gene is a combination of the adaptive value of the gene product, and its expression level [25]. We envisage that equivalently, during the process of *de novo* gene birth, mutational fitness effects can be decomposed into the effect on adaptive value and the effect on expression level. In the model, we use the DFE, together with empirical measurements of mutational effects on expression, to extract a scenario of the evolution of the adaptive value of the *de novo* gene product.

We find that the DFE alone is insufficient to determine the fate of non-genic loci, and the interplay of mutation rate and selection plays a crucial role. We tested the predictions of our model under two mutation rate regimes: In nature, most organisms have a mutation rate in the range 10^−11^-10^−8^ mutations per base-pair per generation (Additional file: Table.1 [18,26–46] – The lower end of mutation rates corresponds to the germline of ciliate *Tetrahymena thermophila* [35] and the higher end corresponds to the plant *Arabidopsis thaliana* [44].). This range falls under the low mutation rate regime. Whereas, population dynamics in laboratory growth competition experiments such as in [47], where multiple variants of a gene compete against each other for a small number of generations, are comparable to the high mutation rate regime.

Our model produced contrasting pictures for these regimes: In the high mutation rate regime, we find that a wide range of biologically reasonable DFEs allow functionalization of non-genic loci. Moreover, this gain of functionality occurred despite the antagonistic effects of locus deletion, particularly for the *C reinhardtii* DFE parameters. In the special case of *de novo* gene birth, mutational effects on adaptive value had a short-tailed distribution, thus implying that the rare, extreme mutations that are characteristic of DFEs are driven solely by mutational effects on expression level.

On the other hand, in the more natural low mutation rate regime, we find that conditions that allow functionalization of non-genic loci are much more stringent, and populations are very sensitive to the effects of locus deletion. Moreover, in the case of *de novo* gene birth, mutational effects on adaptive value had a long-tailed distribution; this implies that in the low mutation rate regime, the rare, extreme mutations in DFEs are driven by both mutational effects on expression level and adaptive value.

However, under both the low and high mutation rate regimes, we find that mutations in adaptive value, rather than expression level are the major drivers for the sustained fitness increase over evolutionary time. Our results can be tested experimentally using high throughput mutational scans on random initial sequences; such experiments stand to offer quantitative insights into the process of *de novo* gene birth.

### Model of non-functional locus adaptation

We set up a population genetic framework to model well-mixed finite populations of fixed size *N*, composed of asexually reproducing, haploid individuals. Fitness of an individual represents growth rate in the exponential phase, which is equivalent to the quantities considered in experiments that measure DFEs (e.g., [18]). In this work, for any individual *i*, we consider the evolution of the fitness contribution *F* (*i*) of a single locus in its genome. We are interested in the probability that the locus persists in the population, and that its fitness contribution increases above some large, predetermined threshold. In this study, we use a fitness threshold of 0.1, which is much larger than the typical effect sizes of incident mutations.

In particular, we examine the special case of *de novo* gene birth, where the functional product of a gene can be either protein or RNA. We decompose the fitness contribution into two quantities: functionality, or *adaptive value* of the expression product (*A*(*i*)), and its *expression level* (*E*(*i*)). Note that this decomposition represents the limiting case where the whole fitness contribution of the *de novo* gene is due to the gene product. It is likely that for emerging genes, the relative contribution of the gene sequence due to its role in gene regulation, or chromosome arrangement is comparable to the contribution of the expression product, and this relative contribution is itself an evolvable feature of the locus. Our definition of fitness of the product is not tied to any specific function, and we assume that *F* (*i*) = *A*(*i*) *× E*(*i*). We note that genes exert their effect on organismal fitness within the context of a molecular interaction network, and never work in isolation [48]. Formally, in our definition, adaptive value *A* represents the *maximal contribution to organismal fitness made by the expression product, conditional upon the underlying cellular molecule interaction network*. Hence, *A* is a simple measure that incorporates the effects of all other genes. In the model, this maximal fitness contribution is realized at the maximal expression level, *E* = 1. And Δ*A* is the change in the maximal fitness contribution of a variant due to a mutation. Notice here that while this decomposition is fairly straightforward for unicellular organisms, the case of multicellular organisms is more complicated: in general, we can expect different cell types to express the same gene at different levels. Therefore, in multicellular organisms, fitness is a function of the expression profile of the gene across all cell types. Hence, we limit our discussion of *de novo* gene birth to unicellular organisms.

The locus of interest is non-genic, with initial adaptive value *A*_0_(*i*) = 0, and an initial expression level *E*_0_(*i*) = 10^−3^ for all individuals. The initial expression level captures leaky expression of intergenic regions [49], which is estimated to be 1000-fold smaller than the level of highly expressed genes [50].

A single time-step is the time it takes for one mutation to occur in the locus. Now, the size of a single locus is very small compared to the whole genome. Hence, it is reasonable to assume that the rest of the genome accrues many more mutations in the time it takes for a single mutation to be incident on our locus of interest. The duration of a time-step for different organisms is equivalent to the inverse of the average mutation rate measured for these organisms: For a locus of ∼100 base pairs, a single model time-step can range between 10^2^ − 10^5^ years for different organisms (Fig.1(B), see also Additional file: Table.1). That is, a single time-step in the model encompasses many generations. Hence populations at different time-steps of the model are non-overlapping; the population at time-step *t* + 1 is composed entirely of the offspring of individuals in the time-step *t* (Fig.1(A)). Offspring incur mutations at each time-step, which affect the locus fitness (Δ*F* (*i*)).

**Figure 1.**
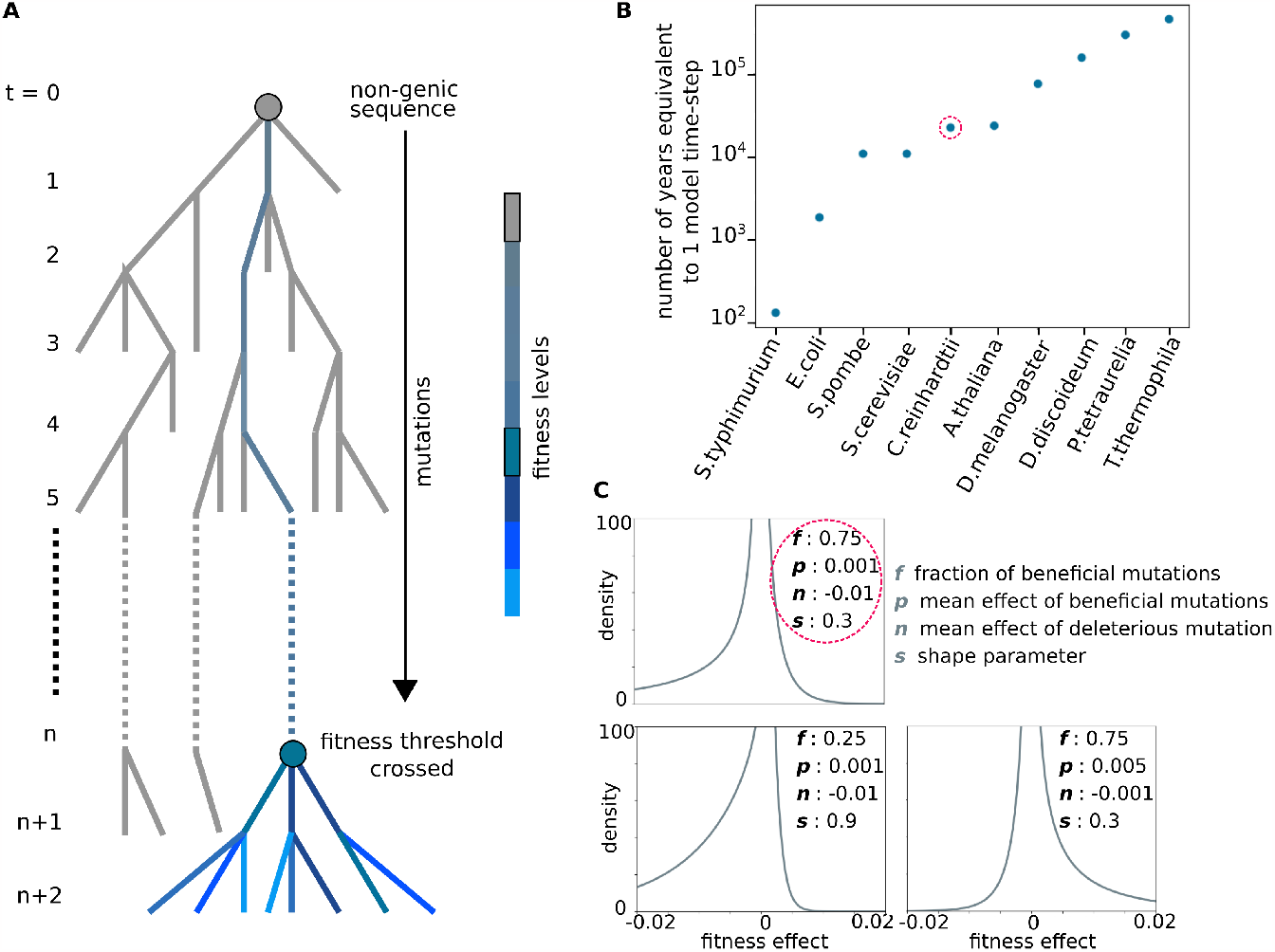
Time-scale and fitness effects of mutations in the model. (A) Phylogenetic tree representing the evolution of a non-genic locus. A model time step *t* spans the average time it takes for a mutation to occur in the locus. The grey dot at *t* = 0 represents the initial non-genic sequence. Grey branches represent lineages that die out, and colored branches represent the lineage that gets fixed in the population. Fitness levels of colored branches in the fixed lineage are indicated in the color bar. The blue dot at *t* = *n* represents the most recent common ancestor of all surviving lineages whose fitness contribution is above the threshold. (B) Estimates of the number of years equivalent to a single time-step of the model in the different species listed on the x-axis. The point representing *Chlamydomonas reinhardtii* is circled in red. See Additional file: Table1 for calculations. (C) Distributions of fitness effects (DFE) for different values of model parameters (listed for each distribution). All DFEs shown here have the same shape parameter, *s* = 0.3, which controls how long-tailed the distributions are. The top left panel represents the DFE with model parameters closest to those measured for *C reinhardtii* in [18]. The bottom left panel represents the DFE with the most deleterious and least beneficial mutations. The bottom right panel represents the DFE with the most beneficial and least deleterious mutations sampled in this work.

In the case of *de novo* gene birth, Δ*F* (*i*) can be decomposed into mutational effects on adaptive value, Δ*A*(*i*), and expression level, Δ*E*(*i*):

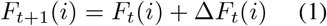

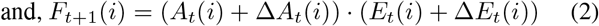

Offspring can also incur structural variations, which in the model leads to the deletion of the locus in that individual. The probability of locus deletion *d* represents the rate of structural variation relative to mutation rate.

In this study, we separately investigate two qualitatively different scenarios:

- When mutation rate is high, and the number of generations between two model time-steps is *<* 2,000, we model the probability that an individual leaves an offspring in the next time-step as being proportional to the fitness *F* (*i*) of the locus.
- When mutation rate is low, and the number of generations between two model time-steps is *>* 2,000, a single variant gets fixed in the population before the next mutation arrives. This is in contrast with the high mutation rate regime, where each model time-step is initialized with multiple variants (Additional file: Section 2). Here, we model the probability of fixation of an individual *i* at each model time-step as being proportional to its fitness *F* (*i*). This increase in fixation probability with fitness is consistent with studies that infer evolutionary outcomes in simpler situations with homogeneous populations and a single mutant [51]. (We describe how we update populations with high and low mutation-rates in Method to update population fitness.)

For simplicity, we do not consider the compounding effects of competition with established genes, and genetic interactions such as linkage and epistasis in the main manuscript (see Additional file: Section 3 for a discussion of evolution in the presence of other non-genic loci and established genes, and the effect of diminishing returns epistasis). Whenever *F* (*i*) ≤ −1, we consider the locus lethal and such individuals cannot produce offspring. We update populations for 1,000 time-steps, equivalent to 0.1-100 million years, depending on the organism and size of the locus (Additional file: Table.1).

Fitness effects of mutations (Δ*F* (*i*)) are drawn from the characteristic DFE of the locus (Fig.1(C)). Multiple studies indicate that long-tails are important features of DFEs, which can be captured by the general form of long-tailed gamma distributions [52, 53]. Therefore, we choose to follow [18], and represent DFEs as two-sided gamma distributions, and characterize them using four parameters: (i) average effect of beneficial mutations *p*, (ii) fraction of beneficial mutations *f*, (iii) average effect of deleterious mutations *n*, and (iv) the shape parameter *s*, where distributions with lower *s* are more long-tailed. In our simulations, we first decide whether a new mutation is beneficial or deleterious by using *f* as the probability of beneficial mutations. We then draw mutational effect sizes: we use a gamma distribution with parameters *p* and *s* for beneficial mutations, and a gamma distribution with parameters *n* and *s* for deleterious mutations (Method to update population fitness).

Mutations in the model represent the mutation types included in [18], which were single-nucleotide mutations and short indels (insertions or deletions of average length ≤ 10 bp) [54]. Note that the quantity reported in experimental studies is the DFE across the whole genome. We account for differences in DFEs across species and locations along the genome by sampling widely across these four parameters *p, f, n, s*. Consistent with the expectation that mutations at non-genic loci should be mostly neutral, the parameters we sample include a large number of DFEs which produce a majority of neutral mutations (Additional file: Section 4).

We also use empirical measurements to estimate the distribution of mutational effects on expression. Studies indicate that mutational effects on expression from established promoters follow a heavy-tailed distribution [55]. More relevant to our study of *de novo* gene birth are the recent measurements of mutational effects on expression from *random* sequences [46], which follow a power law distribution, Pr(Δ*E*) ∼ Δ*E*^−2.25^. We assume in the model that each mutation has two components: a component that affects the adaptive value of the product (Δ*A*(*i*)), and a component that affects expression level (Δ*E*(*i*)). At each time step, we use the above power law distribution to draw Δ*E*(*i*). We then calculate values of mutational effects Δ*A*(*i*) using equations (1) and (2), given distributions of mutational effects on fitness and on expression level (Method to update expression level and adaptive value; see also Additional file: Section 5 for possible deviations from the power-law Δ*E* distribution due to the very small initial values *E*(*i*)).

In all, we survey 432 parameter sets – 108 DFE parameters *p, f, n, s*, and 4 parameter values for *d*, representing the probability of locus deletion – (Fig.1(C)). We simulated replicate populations by performing 100 independent stochastic simulations with the same set of parameters for population sizes *N* = 100, 1,000 (Surveying the space of DFE and locus deletion parameters in populations of various sizes). Simulating replicate populations allows us to estimate the probability that non-genic loci with properties comparable to model parameters can adapt and become functional.

At the end of each simulation, we trace the ancestry of each locus in each individual (Tracing ancestry to find fixed mutations) in order to track the fitness contribution of the *last common ancestor* of the locus: for populations in the high mutation rate regime, the ancestry of all individuals at some time-step *t* can be traced back to a single individual at some previous time-step *t* − *t*_fix_. Whereas, for populations in the low mutation rate regime, all individuals at time-step *t* are the progeny of the single variant that gets fixed in the time-step *t* − 1. We count the number of replicate populations in which the locus is still retained at time-step *t* = 1,000, and the *last common ancestor* of individuals at *t* = 1,000 is fitter than the predetermined fitness threshold of 0.1 (Fig.1(A)).

## Results

### Tuning the number of neutral mutations

Many studies report the abundance of deleterious mutations compared to beneficial mutations, but there is plenty of evidence to the contrary as well [56]. We do not have extensive empirical measurements of the DFE of non-genic loci, nevertheless, since these loci are not expected to be expressed at high levels, it makes intuitive sense to assume that most non-genic mutations should be nearly neutral. In order to reconcile the above factors, we explore a wide range of DFEs around the DFE reported for *C reinhardtii* in [18].

To test the diversity of DFEs we sample in this study, we count the number of positive-effect, negative-effect, and neutral mutations that the different DFEs produce. Theoretically, we expect mutational effects of magnitude 10^−3^ to be neutral for a population of size N = 1000. We use a more stringent condition, and count mutations with effect size *<* 0.5 × 10^−3^ as effectively neutral (we also demonstrate neutrality of these mutations in Additional file: Section 4). Thus, we count mutations of effect ≥ 0.5 × 10^−3^ as positive effect mutations, mutations with effect ≤ −0.5 × 10^−3^ as negative effect mutations, and mutations with effect between −0.5 × 10^−3^ and 0.5 × 10^−3^ as neutral mutations.

In particular, we measured how the parameters of the model control the fractions of positive, neutral, and negative effect mutations (Fig.2(A)), and the magnitudes of positive and negative effect mutations (Fig.2(B)). Importantly, we find that the fraction of neutral mutations decreases with the shape parameter (s).

**Figure 2.**
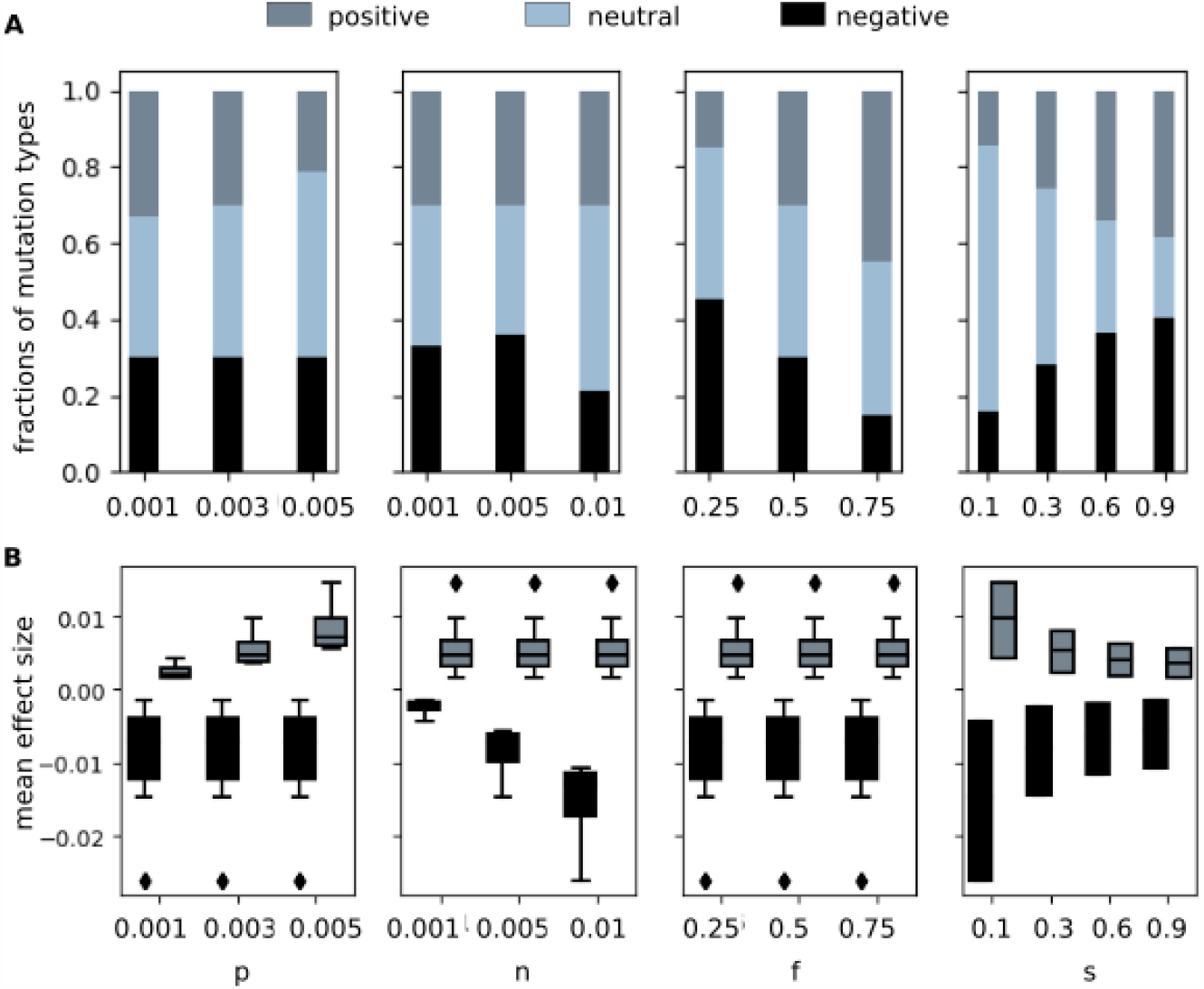
Proportions and magnitudes of positive effect, neutral and negative effect mutations. The plots includes data for 1.08 × 10^9^ mutations: 10^7^ mutations independently drawn using each of the 108 parameters sets (*p, n, f, s*) sampled in this study. In (A,B) grey indicates positive effect mutations, blue indicates neutral mutations and black indicates negative effect mutations. (A) Stacked histograms indicating the average proportions of mutations drawn using parameter values indicated on the x-axis that have positive, neutral or negative effect. (B) Mean effect sizes of positive (grey) and negative (black) mutations drawn using parameter values indicated on the x-axes. Boxes indicate data points, lines within boxes indicate the median, whiskers indicate quartiles, and diamonds indicate outliers.

Out of the 108 DFEs sampled, 50 % of the different DFEs produced more neutral mutations than positive or negative effect mutations. In 52 % DFEs, negative effect mutations were more numerous than positive effect mutations, and in 70 % DFEs, average effect size of negative mutations was larger than that of positive mutations. 18 out of 108 DFEs had more neutral mutations than positive or negative mutations, and produced larger and more numerous negative effect mutations than positive effect mutations. Whereas, only 1 DFE had fewer neutral mutations than either positive or negative mutations, and smaller and fewer negative effect mutations than positive effect mutations. Overall, the parameters we sample cover a broad range of DFEs; many, but not all, of the sampled DFEs match our intuitive beliefs and produce mainly neutral and deleterious mutations.

### Conducive parameters for adaptation

We want to understand which of the 108 DFE parameter sets (*p, n, f, s*) that we test are conducive to adaptation. In order to do this, we simulate the evolution of loci for which incident mutations follow a given DFE in populations of size *N* = 1,000. We perform this simulation in 100 replicate populations for each of the 108 DFEs. These simulations do not include locus deletion (i.e. *d* = 0). We also test how mutation rate influences the fate of non-genic loci: We perform two separate sets of these 10,800 simulations corresponding to population dynamics at high and low mutation rates respectively.

In the model, we keep track of the ancestors of all the individuals at any given time-step. After a sufficient number of time-steps, we find that all individuals at some time-step *t* come from a single common ancestor: In the low mutation rate regime this common ancestor occurs in the immediately previous generation *t* − 1, while in the high mutation rate regime, the common ancestor can be traced back to ∼ 100 time-steps ago. We call this most recent common ancestor of all individuals at time-step *t* the ‘last common ancestor’. Equivalently, we say that the variant of the locus present in this ‘last common ancestor’ is fixed in the population at time *t*. In this analysis, we check whether or not the fitness contribution of the variant in the last common ancestor of all individuals at time-step *t* = 1,000 has crossed the threshold of *F* = 0.1. We call the fraction of replicate populations in which the fitness threshold is crossed the *conducivity* of the corresponding DFE. And we call a given set of DFE parameters *conducive* if its *conducivity* ≥ 0.5; i.e. the fitness of the last common ancestor for individuals at time-step *t* = 1,000 crosses 0.1 in at least 50 out of 100 replicate populations.

#### High mutation rate

##### Rare beneficial mutations are sufficient for adaptation

We find that a majority (80 out of 108) of DFE parameter sets are *conducive* (Fig.3(A) grey bars, see Additional file: FigS8 for *N* = 100). Moreover, the bimodality of the grey histogram in Fig.3(A) indicates that DFE parameters tend to either be highly conducive, or highly repressive to adaptation. As one can anticipate, the conducive DFE parameter sets tend to have high values for the magnitude (*p*) and the frequency of beneficial mutations (*f*), and low values for the magnitude of deleterious mutations (*n*) (Fig.3(A), inset– blue bars, see also Additional file: FigS8(A), inset). In particular, for the DFE parameters closest to *C reinhardtii*, 97% of the N=1,000 replicate populations (52% of N=100 replicate populations) crossed the fitness threshold. Parameters representing *non-conducive* DFEs showed the opposite trends; they tend to have low values for the magnitude (*p*) and the frequency of beneficial mutations (*f*), and high values for the magnitude of deleterious mutations (*n*) (Additional file: FigS9). Non-conducive DFEs are highly repressive, and for 24 out of the 28 *non-conducive* DFEs, none of the *N* = 1,000 replicate population crossed the fitness threshold.

**Figure 3.**
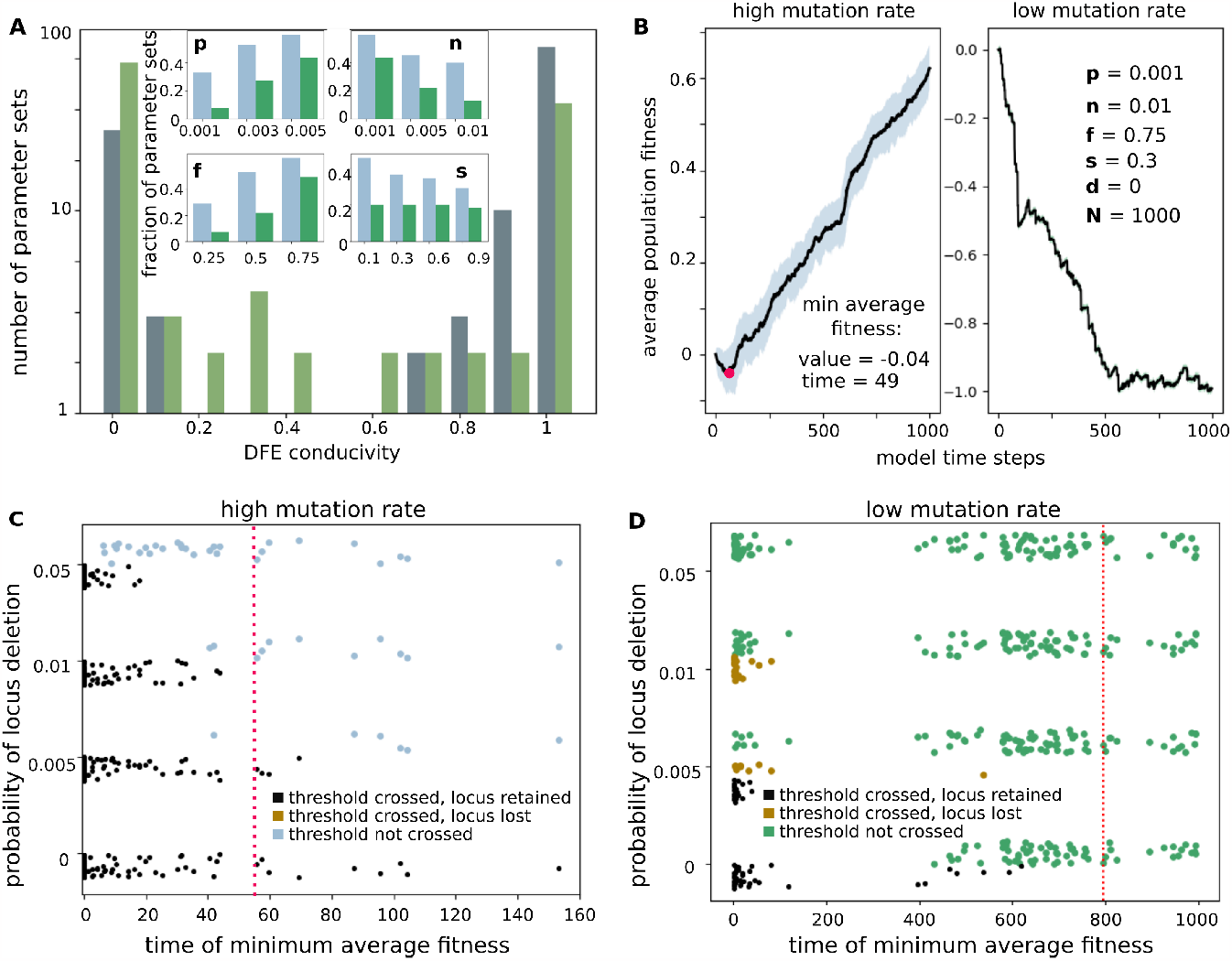
Probability of crossing fitness threshold via accumulating mutations. (A) Histogram for DFE *conducivity* – the fraction of replicate populations (of size *N* = 1,000) with the same DFE where the locus fitness goes on to cross the 0.1 threshold, at the absence of locus deletion (*d* = 0). Bars indicate the number of DFE parameter sets (*p, n, f, s*) in a given *conducivity* bin (Grey bars - high mutation rate regime, dull green bars - low mutation rate regime). This figure includes 108 DFE parameter sets for each of the low and high mutation rate regimes. Inset: Histograms for the fraction of *conducive* parameter sets with given values of parameters *p, n, f*, or *s* (Blue bars - high mutation rate regime and green bars - low mutation rate). (B) Trajectories of population averaged fitness in one of the replicate populations with DFE parameters indicated in the legend, and no locus deletion, *d* = 0, for a population with high (left) and low (right) mutation rates. This DFE is closest to the one reported for *C reinhardtii* in [18]. Black lines show average fitness with shading showing standard deviation. The red point in the left panel indicates the time step at which average fitness reached a minimum value. Each panel includes fitness values for *N* = 1000 individuals over 1000 time-steps. (C,D) Scatter plots showing the effect of locus deletion in 432 parameter sets (4 values of locus deletion rate *d* for each of 108 DFEs). The x-axis indicates the time at which minimum average fitness is achieved, where the average is over all populations with the same DFE parameters, *p, n, f, s* (with random jitter to make point density more apparent). The dotted red line indicates the *time of minimum average fitness* for DFE parameters close to *C reinhardtii*. (C) High mutation rate regime: Black (blue) points represent parameter sets where ≥ 50 % (*<* 50 %) of replicate populations cross the fitness threshold. All populations where the fitness threshold was crossed retain the locus. (D) Black (green) points represent parameter sets where ≥ 50 % (*<* 50 %) of replicate populations cross the fitness threshold. Brown points represent parameter sets where the fitness threshold was crossed in ≥ 50 % of replicate populations, and the locus was subsequently lost in *>* 95 % of the replicate populations.

We also compared the number and magnitudes of positive and negative effect mutations: 21% of DFEs had more numerous and larger positive effect mutations and 100% of these DFEs were *conducive*. However, 45% of the DFEs had more numerous and larger negative effect mutations, and 49% of these very stringent DFEs were also conducive. Among the stringent DFEs, *conducivity* was strongly negatively correlated with the number of negative effect mutations (Pearson’s correlation coefficient *R* = −0.60, p-value = 4.5 × 10^−6^).

In a principal components analysis (PCA) of the proportions and magnitudes of positive, negative and neutral effect mutations, the first three components capture a large part (77.5 %) of the variance in model-generated data (Additional file: Section 6.3). *DFE conducivity* correlates well with all three principal components (PCs). In particular, the first PC is highly correlated with the shape parameter (s) (Additional file: Table2). This is in line with our observation that a substantial number of model loci where positive effect mutations were small and rare could adapt and cross the fitness threshold of 0.1 (Additional file: FigS.6(B)). We find that parameters with low values of the shape parameter (*s*) can compensate for low frequencies, and small average effect size of beneficial mutations (Additional file: FigS.11(A), FigS.12). Notably, while the *conducivity* of DFEs is strongly negatively correlated with the proportion of negative effect mutations (Pearson’s correlation coefficient *R* = −0.68, p-value = 4.9 × 10^−16^), it is positively correlated with both the proportion of positive effect mutations and neutral mutations (R = 0.35, (p-value = 1.8 × 10^−4^), R = 0.3 (p-value = 1.6 × 10^−3^) respectively.

Overall, we find that in the high mutation rate regime, even very rare large-effect beneficial mutations are sufficient for adaptive evolution. We further test our conclusion in an alternative scenario: we use a *non-conducive* DFE (*p* = 0.001, *n* = 0.005, *f* = 0.25, *s* = 0.1), augmented by extremely rare (probability = 0.0001), large effect positive mutations (Δ*F* = 0.15). Consistent with our findings, the evolution of the locus under this scenario leads to adaptation (Additional file: Section 7).

#### Low mutation rate

##### Adaptation requires frequent and large beneficial mutations

In this regime, 43 out of 108 DFE parameter sets are *conducive* (Fig.3(A) dull green bars). While the histogram of DFE *conducivity* is bimodal, there are many more parameters with intermediate *conducivity*. Consistent with the case of high mutation rates, *conducive* DFE parameter sets tend to have high values for the magnitude and the frequency of beneficial mutations (*p* and *f*), and low values for the magnitude of deleterious mutations (*n*) (Fig.3(A), inset-green bars); correspondingly, parameters representing *non-conducive* DFEs tend to have low values for the magnitude (*p*) and the frequency of beneficial mutations (*f*), and high values for the magnitude of deleterious mutations (*n*) (Additional file: FigS9).

However, in the low mutation rate regime, the requirements for gene birth are much more strict, and for the DFE parameters closest to *C reinhardtii*, none of the replicate populations crossed the fitness threshold (Fig.3(B), right panel). Overall, 24% of DFEs that had more numerous and larger positive effect mutations were *conducive*, and only 1.5% of the very stringent DFEs that had more numerous and larger negative effect mutation were *conducive*.

Moreover, in the principal components analysis, DFE *conducivity* correlates well with the second and third, but not the first PC (Additional file: Table2). In line with this, the shape parameter (*s*), which is highly correlated with the first PC, does not compensate for small and low frequency beneficial mutations (Additional file: Table2, FigS11(B)). Consistently, the *conducivity* of DFEs is strongly negatively correlated with the proportion of negative effect mutations (Pearson’s correlation coefficient *R* = −0.58, p-value = 4.8 × 10^−11^) and positively correlated with the proportion of positive effect mutations (*R* = 0.50, p-value = 3.3 × 10^−8^), but shows weak and non-significant correlations with the number of neutral mutations (*R* = 0.1, p-value = 0.3). Thus in contrast with the high mutation rate regime, rare beneficial mutations are not sufficient to drive adaptation in the low mutation rate regime.

### Population dynamics and fitness trajectories

We next looked at the trajectories of population averaged fitness in order to find out whether there are significant regularities in the dynamics of adaptation.

#### High mutation rate

##### Fitness trajectories transition from a mutation dominated to a selection dominated phase

We find that the fitness trajectories of populations where the fitness threshold is crossed have a typical shape: First the average population fitness decreases to reach a minimum value. Beyond this, the average population fitness increases roughly linearly (Fig.3(B), left panel). We intuit that these two phases can be explained by the opposing effects of new incident mutations and of selection: In the first phase where the population average fitness decreases, fitness is likely dominated by the effects of new mutations which are more likely to be deleterious/neutral (In 52% DFEs, negative effect mutations were more numerous than positive effect mutations, and in 70% DFEs, average effect size of negative mutations was larger than that of positive mutations). In the second phase, population average fitness increases because the effects of selection become apparent after a sufficient amount of time has passed, and dominate over the effect of new, incoming mutations.

These fitness trajectories are reminiscent of the dynamics of learning through adaptive strategies in gambling problems, where an initial phase of loss of capital due to the cost of learning is followed by recovery [57]. Although, at the level of a single organism, the fitness effects of new mutations are independent of the history of mutations that came before, natural selection endows populations with a form of memory [58, 59].

Two numbers indicate the point in the trajectory at which selection leads to consistent improvement in fitness: *minimum average fitness* and *time at which minimum fitness is achieved* (Fig.3(B), left panel). As expected, populations with lower minimum fitness spend longer in the decreasing fitness regime (Pearson correlation coefficient between *minimum fitness* and *time of minimum fitness* = -0.87, with p-value = 5.5 × 10^−34^. See supplementary figure FigS.9). We interpret the *time of minimum fitness* as representing the amount of time it takes for the effect of selection to start dominating over the effect of new incident mutations. In the high mutation regime, DFE parameters, especially *p* and *f*, are significantly correlated with time of minimum fitness (Pearson’s *R* = −0.41 (p-value = 9.3 × 10^−6^) and −0.52 (p-value = 1.9 × 10^−8^), respectively. See Additional file: Table3)).

#### Low mutation rate

##### Fitness trajectories always governed by incident mutations

In simulations with low mutation rates, a single variant gets fixed in the population at each model time-step. Therefore, although the absolute values of fitness vary across model time-steps, the distribution of relative fitness remains unchanged, and are governed by the fitness effects of incident mutations. Hence, in the low mutation rate regime, ‘learning’ occurs during the many rounds of selection in the generations intervening two mutational time-steps, and is not visible at the time-scale of model time-steps. Consistently, even for *conducive* DFE parameters, fitness trajectories do not have a selection dominated phase of persistent fitness increase (Additional file: Section 9).

### Dynamics of adaptation, locus loss and retention

We next test the effect of large structural variations that can delete the entire locus in an individual on the chances that the locus can adapt and become fixed in the population. Note that we are concerned with the general case of gain of functionality by a locus, which may or may not be due to expression of a product (protein or RNA). In the particular case where the locus expresses a protein product, there are many ways to erase functionality, such as by disrupting mRNA export from the nucleus, ribosome binding site, or truncating gene products through non-sense mutations. Here, we only study the effect of locus deletion, which affects all loci irrespective of their particular mode of function.

#### High mutation rate

##### Mutations drive adaptation despite the effect of locus deletion

When *d >* 0, The effect of locus deletion can be understood in terms of a competition between two sub-populations: the sub-population that has lost the locus, and therefore lacks any fitness contribution from it, and the sub-population that retains it (Additional file: Section 10). The probability that the sub-population that has lost the locus takes over increases with *time of minimum fitness* as calculated for the case where *d* = 0: the longer the average fitness remains negative, the more probable is the loss of the locus from the whole population. Therefore, fewer replicate populations with DFEs such that minimum fitness is reached later go on to cross the fitness threshold of 0.1 (Fig. 3(C), Additional file: FigS.13(B)). As a consequence, the number of *conducive* parameter sets (out of 108 sets overall) reduces from 80 at *d* = 0, to 74 at the plausible value of *d* = 0.005, and to 69 and 50 at the inordinately high values of *d* = 0.01 and 0.05, respectively. Particularly, the DFE closest to *C reinhardtii*, for which *minimum fitness* and *time of minimum fitness* averaged across all replicate populations are −0.035 and 55.7 respectively, is still *conducive* at *d* = 0.005 (Fig.3(C), red dotted line).

#### Low mutation rate

##### Structural variations lead to locus loss despite adaptation

In this regime, a single variant gets fixed at every time step, which results in populations being very sensitive to locus deletion. The dynamics can still be understood in terms of sub-populations, which now compete for fixation during the many intervening generations within one model time step: At every time step, the locus gets deleted in a small fraction of the population, and there is a finite probability that an individual from this sub-population takes over and becomes fixed in the population. This probability is proportional to the relative fitness of the sub-population that has lost the locus (Additional file: Section 2).

Hence, adaptation in this regime requires consistent fixation in successive time-steps of variants from the sub-populations that contain the locus. Additionally, the larger the magnitude of beneficial mutations that are fixed in subsequent time-steps, the faster the decrease in relative fitness of the sub-population that has lost the locus. Consistent with this picture, many populations where the fitness of the locus does not grow fast enough lose the locus despite having crossed the threshold of 0.1 (Fig.3(D), brown points): For the plausible value of *d* = 0.005, for 29 DFE parameter sets *>* 50% replicate populations cross the fitness threshold of 0.1; and for 9 of these 29 parameter sets the locus is subsequently lost from *>* 95% of the replicate populations. This result is in agreement with the observations in

*Drosophila*, where proto-genes that get fixed in populations are very often lost [60] despite showing signs of adaptation [61].

For *d* = 0.005, all 29 DFE parameters for which the locus is retained (Fig.3(D), black points) have more numerous positive effect mutations than negative effect mutations, and 75% of these have larger positive effect mutations than negative effect mutations. For the high value of *d* = 0.01, 20 DFE parameter sets are *conducive* to adaptation, and the locus is subsequently lost from *>* 95% of the replicate populations for all of these parameter sets. For *d* = 0.05, there are no *conducive* DFE parameter sets.

### Functionality and expression as drivers of adaptation

In the model, we draw mutational fitness effect (Δ*F*), and mutational effect on expression level (Δ*E*) independently from their respective distributions. We then derive Δ*A* using the relation in equation (2). Note that the distribution of Δ*A* we report is a population measure which depends on the standing fitness variation. Hence, we expect the distinct population dynamics in the high and low mutation rate regimes to lead to distinct distributions of Δ*A*. The shape of the distribution of Δ*A* that we derive here is an important indicator, and reflects how single small mutations affect the functionality of newly emerging genes.

We also ask whether fitness increase during *de novo* gene birth is driven more by changes in adaptive value or expression level. As a measure of the strength of driving, we use correlations between the trajectory of population averaged fitness and the trajectories of population averaged adaptive value and expression level. (Although instantaneous values of fitness, expression level and adaptive value have a simple relationship (*F* = *A × E*), the trajectories of these quantities over evolutionary time also include the effect of selection and heritability, and their correlations are therefore non-trivial.) To illustrate, although Δ*F* (*i*) and Δ*E*(*i*) for mutations are drawn from independent distributions, the process of selection effectively links fitness and expression level, and imposes correlations between their evolutionary trajectories (Additional file: compare FigS.19(A), FigS.20). The correlations between the trajectories of fitness, expression level and adaptive value measure the degree to which the direction of changes in average fitness levels (increase/decrease) reflect the directions of change in average expression levels and adaptive values in the population.

#### High mutation rate

##### Functionality drives sustained adaptation, while expression drives extreme mutational events

In this regime, our decomposition of fitness into expression level and adaptive value yields an exponential distribution for mutational effects on adaptive value (Fig.4(A), Additional file: Section 12). The short-tailed nature of Δ*A* distribution implies that mutations with large effects on adaptive value are highly unlikely. Therefore, the extreme mutational effects on fitness, which underlie the long tails of DFEs, are likely to be due to large changes in expression level. We anticipate that our result on the Δ*A* distribution can be measured experimentally; for instance, one could generate random mutants of known proto-genes, and measure the fitness effects of these mutations using techniques demonstrated in [16]. We also expect that dynamics in the high mutation rate regime should be more suited to capture conditions in laboratory experiments (as in [47]).

**Figure 4.**
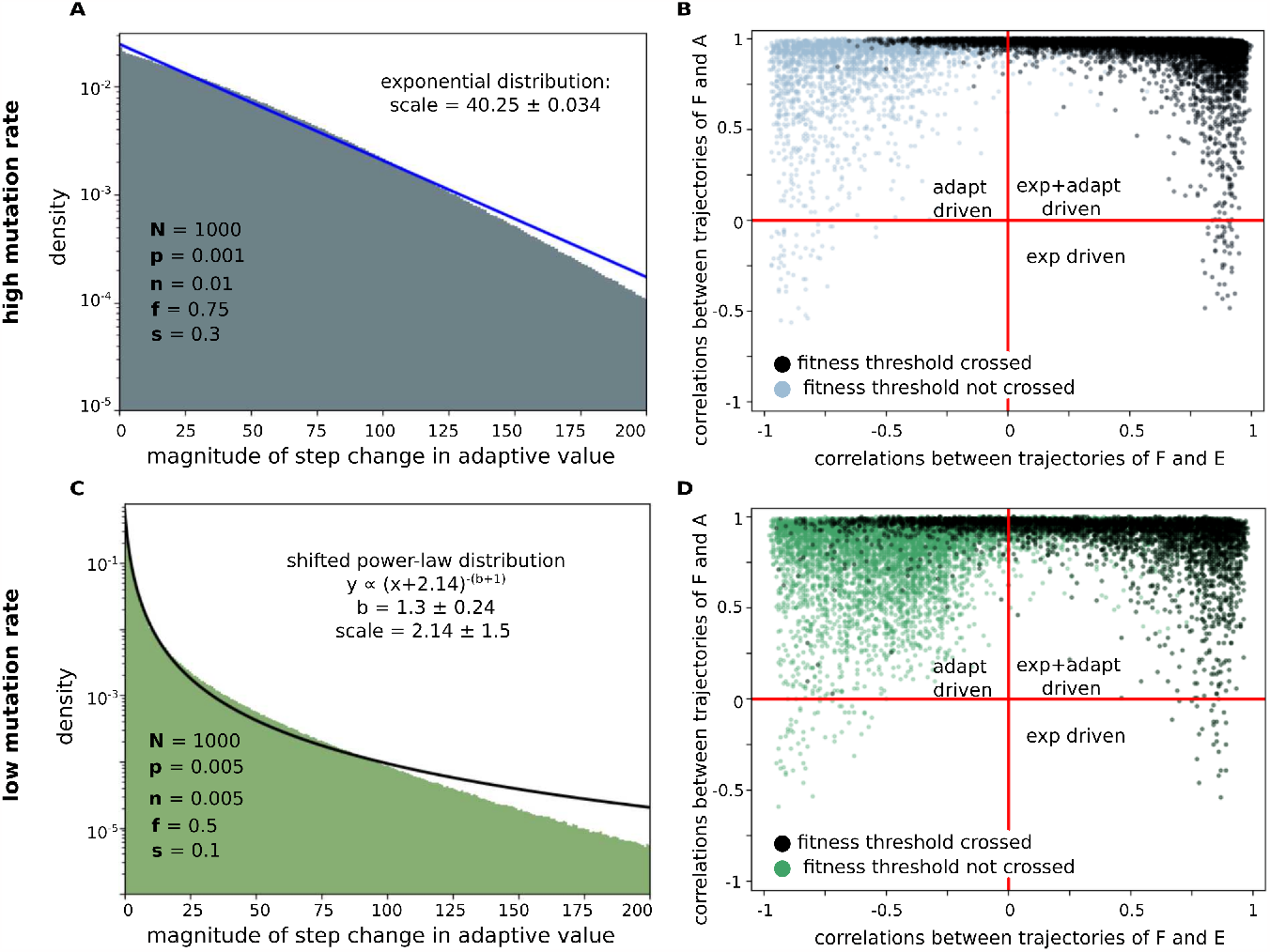
Distribution of Δ*A* and the drivers of fitness increase. We used the distribution reported in [46] to generate Δ*E* in order to obtain trajectories of expression level, *E*, and adaptive value, *A*, from which we infer values of Δ*A*. (A,C) Histograms of 10^8^ Δ*A* values each (pooled across all 1000 individuals in all 100 replicate populations for 1000 time-steps) with (A) High mutation rate dynamics and *Chlamydomonas* DFE parameters [*p, n, f, s*] = [0.001, *-* 0.01, 0.75, 0.3, 0] with an exponential fit (blue). (C) Low mutation rate dynamics and the most stringent *conducive* DFE parameters [*p, n, f, s*] = [0.005, − 0.005, 0.5, 0.1], with a fit to a power-law distribution (black). (B,D) Scatter plots of correlation of population-averaged fitness trajectories with trajectories of population-averaged expression level (x-axis) and trajectories of population-averaged adaptive value (y-axis), in populations following high (B) and low (D) mutation rate dynamics. The black (blue/green) points represent populations that cross (do not cross) the fitness threshold. Overall, each plot contains 108 ×100 points representing all replicate populations across all parameter sets for *N* = 1,000 populations. Red lines demarcate regions where fitness change is driven by changes in expression level (bottom right), driven by changes in adaptive value (top left), or by both expression level and adaptive value (top right). As expected, replicates that cross the threshold (black points) are absent from the bottom left region, where trajectories of both adaptive value and expression level are negatively correlated with the fitness trajectory.

At the same time, correlations between the trajectories of fitness, expression level and adaptive value indicate that increase in fitness was driven more by the adaptive value than by expression level: the distribution of Pearson’s correlation coefficients for adaptive value is sharply peaked at 1, whereas that of expression level is spread broadly (Fig.4(B), Additional file: FigS.19).

Overall, we find that in the high mutation rate regime, sustained adaptation during gene birth is driven more by the product’s adaptive value rather than its expression level. While extreme mutational events, which become important in facilitating adaptation when most beneficial mutations are small and infrequent (i.e. small *f* and *p*), are more likely to be due to changes in expression level.

#### Low mutation rate

*Expression and adaptive value both drive extreme mutational events* In this regime, our decomposition of fitness into expression level and adaptive value yields a heavy-tailed power-law distribution for mutational effects on adaptive value (i.e. a distribution with a power-law tail: Fig.4(C)). Therefore, in contrast to the high mutation rate regime, extreme mutational effects on fitness are driven by large changes in both expression level and adaptive value. However, correlations between the trajectories of fitness, expression level and adaptive value indicate that similar to the high mutation rate regime, increase in fitness was driven more by the adaptive value than by expression level (Fig.4(D)).

## Discussion

A majority of studies in genomics and genetics are concerned with the function and evolutionary course of known genes and their regulation. Recent discoveries have shifted our focus towards the evolution of non-genic loci; particularly, experimental studies that demonstrate the adaptive potential of random sequences [62–65]. Furthermore, genomics studies that indicate the frequent occurrence of *de novo* gene birth demonstrate a need for general, theoretical investigations of the evolution of non-genic loci [11]. In this work, we attempt to describe the process of functionalization of non-genic genomic loci in a simple population genetic model. In terms of the Pittsburg model proposed in [2], we investigate the process by which a genomic locus with physiological implications (contribution to organismal fitness) transforms into a locus with evolutionary implications. We explore the space of biologically reasonable effects of spontaneous mutations and identify features of genomic loci that are conducive to functionalization. We also demonstrate how the outcomes of non-genic evolution depend on mutation rate.

Our results present a contrasting picture for organisms with high mutation rates versus those with low mutation rates: In organisms with high mutation rates, our model predicts that a wide range of parameters that govern mutational fitness effects (DFE) are conducive to locus functionalization. Conducive DFEs include many conservative DFEs, where positive effect mutations are small and few. Adaptation in this regime was also robust to the antagonistic effects of structural variation that leads to locus deletion.

On the other hand, mutation rates for most organisms fall in the range 10^−11^-10^−8^ per base-pair per generation, and are thus in the low mutation rate regime. In this regime, conditions for locus functionalization were much more stringent. Importantly, populations in this regime are sensitive to structural variations that cause locus deletion, and even adaptive loci are often lost. Thus, our model supports the view that gain for functionality by most non-genic sequences should be rare. This is in congruence with the observation that most *de novo* genes are born in regions adjacent to established genes, where sequences have an increased chance of expression though read-through [66], and perhaps also a lower probability of deletion due to linkage.

In the special case where non-genic adaptation leads to *de novo* gene emergence, we use a simple relation to resolve the fitness contribution of *de novo* genes into its adaptive value and expression level. Using the distributions for fitness and expression level, we could infer the distribution of adaptive value, which is difficult to measure directly. We envisage that such a method, where information about measurable quantities can be used to infer constraints on distributions of quantities that are difficult to measure, can be extrapolated to get a deeper view into the mechanism of *de novo* gene birth: Expression level can be further broken down into various molecular mechanisms such as the accessibility and affinity of the locus to polymerases, and the adaptive value can be expressed as a composite of the stability, foldability, and interactions of expression products.

Our results also suggests that the process of adaptation is likely to be different for *de novo* genes and established genes: We find that in the case of *de novo* gene birth, the increase in fitness was driven more by the adaptive value than by expression level in both the low and the high mutation rate regimes. This effect is likely to be a special feature of *de novo* gene birth, where initially both adaptive value and expression levels are very low. Whereas in the case of established genes, evolution of expression level is known to play a role in adaptation [67–69].

In this study, we present a minimal framework to study non-genic adaptation. Below, we discuss some important limitations of this framework, and suggest directions in which the model can be expanded which are particularly relevant to the study of *de novo* genes:

- **Initial conditions:** In our study, the locus of interest initially has no adaptive value, and expression level corresponding to pervasive, leaky expression. However, induced expression studies on random sequences indicate that inter-genic loci can be expected to have non-zero adaptive value [65]. Additionally, some non-genic regions where *de novo* genes have been found, such as 3’ ends of established genes, have higher expression levels than others [66]. These results warrant a wider sampling of initial values of adaptive values and expression levels.
- **RNA versus protein *de novo* genes:** A *de novo* gene in our model can represent either RNA or protein gene, and both protein and RNA products of genes can contribute to organismal fitness [70]. However, protein and RNA gene emergence are expected to be different: (a) Sequence requirements for translation are more stringent than the requirements for transcription; a protein product necessarily requires a ribosome binding site, start and stop codons, and the correct reading frame. This makes the probability that a locus first starts expressing a protein lower, and increases the probability that mutations can destroy the product. Therefore, the forms of Δ*E* and deletion probability *d* are likely to be different for proteins versus RNA. (b) ‘Enabler mutations’ that allow a non-coding region to first start producing a protein product can be expected to have a large effect on fitness. Thus, improved models should consider protein and RNA gene emergence separately. This limitation also points to empirical measurements that could aid in the formulation of improved theoretical studies: For example, for *de novo* genes whose enabler mutations are known (e.g. [9, 71]), fitness effects of these enabler mutations can be experimentally measured.
- **Fitness contribution of DNA sequence versus expression product:** Our decomposition of fitness contribution into adaptive value (*A*(*i*)), and expression level (*E*(*i*)) of the gene product represents the limiting case where the whole fitness contribution of the *de novo* gene is due to the gene product. However, especially for emerging genes, it is likely that a non-negligible part of the fitness contribution comes from the DNA sequence itself due to its role in gene regulation, or chromosome organization. While current experimental efforts are focused on the molecular basis of functionality of *de novo* gene products (such as in [16, 66]), delineating the relative contribution to fitness by the gene product versus the DNA sequence constitutes an important direction for future experiments. It would also be important to understand how this relative contribution of expression product versus gene sequence evolves during gene birth. In future versions of the model, this aspect can be incorporated in the form of a variable *r* ∈ [0, 1], such that

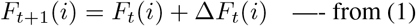

and,

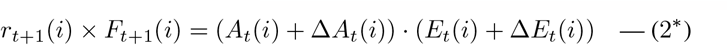 Where *r* represents the fraction of the fitness contribution that can be attributed to the gene product. In the current model, we explore a space where *r* = 1.
- **Genetic interactions:** For simplicity, we consider genomic loci as independent, and do not include genetic interactions such as linkage and epistasis. However, many *de novo* genes are found in the vicinity of well-established genes [8]. Moreover, evolutionary trajectories of loci that are genetically linked to strongly selected, well-established genes can deviate significantly from our model prediction, and hence warrant a separate treatment [72]. In future studies, one could extend the present model to investigate the evolutionary trajectories of two or more interacting loci, some of which represent established genes and have high initial fitness contribution (See Additional file: Section 3 for a discussion of a multi-locus models). Alternatively, our model can also be used to represent mutation scan experiments, such as in [46] where the genomic background is kept constant. In this case, the time-steps in the model represent rounds of experiments involving mutagenesis and artificial selection.
- **Multicellular organisms:** The same gene can have different expression levels in different cell-types of multicellular organisms, thus the adaptive value of a gene is now a function of its expression profile across different cell-types. Therefore, the decomposition of fitness contribution of a gene into its expression level and adaptive value is not straightforward for multicellular organisms.
- **Other important processes:** In this work, we study the role of mutations caused by small indels and polymorphisms, for which quantitative DFE measurements are available. However, evidence also indicates a role for other molecular processes such as transposition, in *de novo* gene birth [60]. Thus, a more complete model would include fitness effects of transposition; although transposition is a less frequent event than indels/ polymorphisms, and we do not have extensive quantitative measurements of fitness effects.
- **DFE sampling:** We sample a broad range of DFEs around an experimentally measured distribution of mutational fitness effects, which makes it likely that we cover the space of real DFEs. However, the distribution of real DFEs across genomic loci and different organisms is not well-known, and our sampling of DFEs is not expected to be identical to the spread of real DFEs.

In addition, the generality of our results is likely to be limited due to dearth of relevant data. Most importantly, we use experimental measurements of DFE and mutational effects on expression that are taken from different organisms: in different organisms, distinct mechanisms produce mutations, therefore the frequencies of different mutation types and its effects may vary. Although, the long-tailed nature of DFEs [52, 53] and mutational effects on expression [46, 55] have both been observed in independent studies, measurements performed in the same organism could provide important details, for instance the correlations between the effect of a mutation on expression and on fitness. Secondly, the DFE of loci remain constant in our model, while mutational fitness effects are known to vary over evolutionary time due to various causes, such as change of environment or diminishing returns epistasis [73–75]. An extended model that includes a consideration of DFE variability would provide valuable insight into the robustness of our results (Additional file: Section 3.3).

We anticipate that our results can be tested and the shortcomings of our model can be addressed through experiments, especially mutational scans such as those in [46]: For example, one could design experiments that monitor the fitness effects of mutations on random sequences and concomitantly detect expression from these random sequences. Alternatively, the evolution of adaptive value of expression products can be directly examined in experiments where random sequences are placed under constitutive, high expression promoters (such as in [62,65]); in this case the fitness effects of mutations directly correspond to the adaptive value of the product. These experiments, together with theoretical approaches like ours, provide us with means to test and compare the adaptive potential of non-functional genomic sequences, and the general mechanisms of *de novo* gene birth across various organisms.

## Conclusions

Our study represents the first population model to analyse the mechanism of *de novo* gene birth within the context of non-genic adaptation. Specifically, we characterize non-genic sequences in terms of their DFE: an empirically measurable quantity. We demonstrate how the DFE, a property of the genomic locus, and mutation rate, an important population dynamical quantity, interplay to influence the fixation of emerging genes. In its current minimal form, our model is already able to capture intriguing phenomena, such as the loss of adaptive proto-genes [60, 61]. We discuss how our model can be expanded to include essential features relevant to *de novo* gene birth, such as the transcriptional piggy-backing of emerging genes on nearby established genes [66], as well as the importance of untangling the fitness contributions of the DNA sequence, RNA product and the protein product of newly emerging genes. Our framework can be used to explore a rich space of processes relevant to non-genic adaptation, and in conjunction with experiments such as mutational scans of intergenic sequences, it can significantly contribute to the discovery of mechanisms of *de novo* gene birth.

## Methods

### Surveying the space of DFE and locus deletion parameters in populations of various sizes

We look at populations of sizes *N* = 1000, and scan across DFEs with parameters *p* = [0.001, 0.003, 0.005], *f* = [0.25, 0.5, 0.75], *n* = [0.001, 0.005, 0.01] and *s* = [0.1, 0.3, 0.6, 0.9]. We look at locus deletion probabilities *d* = [0, 0.005, 0.01, 0.05]. We perform simulations for each parameter set in both high- and low-mutation rate regimes.

In addition, we also look at populations of size *N* = 100, for which we scan across parameters *p* = [0.001, 0.003, 0.005], *f* = [0.25, 0.5, 0.75], *n* = [0.001, 0.005, 0.01], *s* = [0.3, 0.6, 0.9] and *d* = [0, 0.005, 0.01, 0.05]. For *N* = 100 populations, we test only the high mutation rate regime.

For each parameter set, we simulate 100 replicate systems. In all, we look at 118,800 systems: 86,400 *N* = 1,000 systems, and 32,400 *N* = 100 systems. All codes used to generate and analyze data are written in Python3.6.

### Relative fitness of individuals

For a population of size *N*, fitness of individuals at time-step *t* are stored in the real-valued vector *F*_*t*_ of length *N*, where the fitness of any individual *i* is *F*_*t*_(*i*). We also keep track of the individuals that have lost the locus due to deletion in the binary vector *L*_*t*_ of length *N*, such that *L*_*t*_(*i*) = 1 implies that individual *i* contains the locus at time-step *t*, and *L*_*t*_(*i*) = 0 implies individual *i* has lost the locus. Here, *L*_*t*_(*i*) = 0 automatically implies *F*_*t*_(*i*) = 0.

In the model, only individuals with fitness *>* −1 are viable, and capable of producing progeny. And individuals in the current population that produce progeny are selected on the basis of their normalized relative fitness.

Let minfit_*t*_ be the minimum fitness among viable individuals in *F*_*t*_.

We define allfit_*t*_ 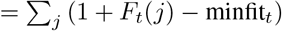, for *j* such that *F*_*t*_(*j*) *>* −1. The normalized relative fitness of individuals is then given by the binary vector relfit_*t*_ of length *N*, where

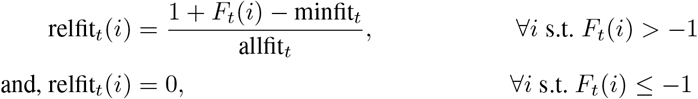

Therefore, even if *F*_*t*_(*i*) = 0, relfit_*t*_(*i*) can be non-zero if minfit_*t*_ *<* 0.

### Mutation dynamics

Progeny of the current population incur mutations. The mutation effects are drawn from 2-sided gamma distributions governed by the parameters *p* (average effect of beneficial mutations), *f* (fraction of beneficial mutations), *n* (average effect of deleterious mutations), and *s* (shape parameter). The values of fitness effects of mutations incurred by each individual at time-step *t* is stored in a real valued vector mut_*t*_ of length *N*, where

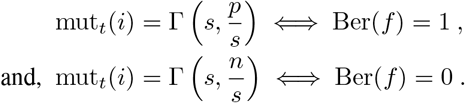

Here Γ (*κ, θ*) represents a number drawn from the gamma distribution with shape parameter *κ* and scale parameter *θ*, and Ber(*f*) is the Bernoulli random variable which equals 1 with probability *f*.

Progeny can also lose the locus with probability *d*. Thus, the updated fitness levels of the population is given by *F*_*t*+1_(*i*) = 0, if *Anc*_*t*+1_(*i*) did not contain the locus, or if the individual loses the locus in the current time step. Otherwise, *F*_*t*+1_(*i*) = *F*_*t*_ (Anc_*t*+1_(*i*)) + mut_*t*_(*i*).

### Method to update population fitness

In the model, the dynamics of selection and mutations interplay and effect how variants of the locus are inherited across model time-steps. We keep track of the ancestry of each individual across time-steps. We do this in order to trace back the *last common ancestor*. The *conducivity* of DFE parameters in the model depends upon whether or not the fitness contribution of the variant of the locus in the last common ancestor of individuals at *t* = 1,000 crosses the threshold of 0.1. Let Anc_*t*+1_ ∈ N*NX*1 be the list of individuals chosen from the current time-step *t* to leave progeny. In other words, Anc_*t*+1_ is the list of ancestors of the population at time-step *t* + 1.

- For populations with high mutation rate **(<2**,**000 generations between mutational time-steps**): Pr(Anc_*t*+1_(*j*) = *i*) ∝ relfit_*t*_(*i*), *∀i, j* ≤ *N*. That is, more than one individual from time-step *t* leaves offspring in time-step *t* + 1.
- For populations with low mutation rate (**>2**,**000 generations between mutational time-steps**): a single variant gets fixed in the population. That is, Anc_*t*+1_ contains a single individual. The probability that an individual *i* is fixed in the population during the intervening generations between two mutational time-steps is proportional to relative fitness relfit_*t*_(*i*) (see Additional file: Section 2).

### Method to update expression level and adaptive value

In the model, we assume *F* (*i*) = *A*(*i*) *× E*(*i*) for any individual *i*. For a population of size *N*, expression levels of the locus at time-step *t* are stored the real valued vector *E*_*t*_ of length *N*, where the expression level of some individual *i* is *E*_*t*_(*i*). For an individual that has lost the locus due to deletion, *L*_*t*_(*i*) = 0, which automatically implies *E*_*t*_(*i*) = 0.

Initially, the expression level of the locus across the population is distributed around 0.001, and reflects leaky expression. At each time step, a mutation is incurred in the locus of interest by each individual in the population. The effect of each mutation has two independent components: one that effects expression level (Δ*E*) and one that effects adaptive value (Δ*A*). Therefore, at each time step, the expression levels and adaptive values across the population change as individuals are selected and their progeny incur mutations.

The effect of mutations on expression level incurred by each individual at time-step *t* is stored in real valued vector Δ*E*_*t*_ of length *N*. The magnitude of Δ*E*_*t*_(*i*) are drawn from a power law distribution such that *Pr*(|Δ*E*_*t*_(*i*)| = *x*) = *x*^−2.25^ for *x* ≥ 0. We assume that a Δ*E*_*t*_(*i*) is negative with probability 0.5.

The updated expression levels of the population are therefore given by *E*_*t*+1_(*i*) = 0, if *Anc*_*t*+1_(*i*) did not contain the locus, or if the individual loses the locus in the current time step. If the individual does contain the locus, *E*_*t*+1_(*i*) = *E*_*t*_ (Anc_*t*+1_(*i*)) + Δ*E*_*t*_(*i*).

Note that the values of expression level in the model are bounded within [0.001, 1] corresponding to leaky expression and maximal possible expression respectively. In the simulation, whenever *E*_*t*+1_(*i*) *<* 0.001 or *E*_*t*+1_(*i*) *>* 1, we reset it to 0.001 and 1, respectively. Since the initial expression levels are very low, *E*_*t*+1_(*i*) never crossed 1 in any simulation. In a run of 1000 time steps, *E*_*t*+1_(*i*) crosses 0.001 on average 40 times (Additional file: FigS7).

We then calculate the corresponding changes in the adaptive value of the locus at each time step: *A*_*t*_(*i*) = *F*_*t*_(*i*)*/E*_*t*_(*i*). From this, we can calculate the change in adaptive value due to mutation as Δ*A*_*t*_(*i*) = *A*_*t*+1_(*i*) − *A*_*t*_ (*Anc*_*t*+1_(*i*)).

### Tracing ancestry to find fixed mutations

This method allows us to identify the mutant that gets fixed in the population at time-step *t* = 1000. In the model, we count and report the number of replicate populations in which the fitness contribution of this fixed mutant crosses the threshold of 0.1.

For populations with **high mutation rate**: In order to find the fitness value of the mutant that gets fixed in the population at time-step *t*, we start with the list of ancestors of individuals Anc_*t*_ at time-step *t*.

Let *X*_*t*_ = *{i, ∀i* ∈ Anc_*t*_*}* be the set of unique ancestor identities. We then recursively find *X*_*t*−*n*_ = *{i, ∀i* ∈ *{*Anc_*t*−*n*_(*j*), *∀j* ∈ *X*_*t*−*n*+1_*}}* as the set of unique ancestor identities for *n* = 1, 2, 3…*t*_0_, where 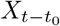 is the first singleton set encountered.

This set contains the individual which occurred at time-step *t* − *t*_0_ − 1, whose locus variant is inherited by every individual at time-step *t*. The fitness value of the mutant fixed in the population at time-step *t* is then 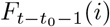, where 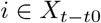.

For populations with **low mutation rate**: the fitness value of the mutant that gets fixed in the population at time-step *t* is simply Anc_*t*_.

## Abbreviations

DFE: Distribution of fitness effects

## Declarations

## Acknowledgements

We thank John McBride, Luca Peliti and Anne-Ruxandra Carvunis for very helpful discussions. We also thank anonymous reviewers of the manuscript for their deep evaluation of the model and its results.

## Funding

This work was supported by the Institute for Basic Science (Grant IBS-R020-D1).

## Availability of data and materials

All data generated or analysed during this study are included in this published article, its supplementary information files and publicly available repositories. Codes and data generated in this study are available at the Dryad repository: mani, somya (2023), data for ‘Gene Birth in a Model of Non-genic Adaptation’, Dryad, Dataset, https://doi.org/10.5061/dryad.fbg79cnxx

## Ethics approval and consent to participate

Not applicable.

## Competing interests

The authors declare that they have no competing interests.

## Consent for publication

Not applicable.

## Authors’ contributions

SM: conceived the project, designed research, developed models, performed simulations, analysed data and wrote the manuscript. TT: conceived the project, supervised research, and wrote the manuscript. All authors read and approved the final manuscript.

## Additional file: Figures S1-S21, Table 1-3

**Table 1**- Estimates of one model time-step in various species **Figure S1**- The distribution of relative fitnesses and absolute fitness values found to fix in the detailed simulation is statistically similar to the distribution of relative fitnesses and absolute fitness values of individuals simply picked with a probability proportional to their relative fitness. **Figure S2**- Comparison of adaptation of loci in the main model versus the multi-locus model. **Figure S3**- Comparison of adaptation of loci in the main model versus a model where *de novo* gene evolution occurs in the presence of an independently evolving established gene. **Figure S4**- Trajectories of population averaged fitness subject to diminishing returns epistasis. **Figure S5**- Neutrality of mutants with fitness effect = 0.5 × 10^−3^ in N=1,000 populations. **Figure S6**- Fraction of positive effect mutations (fitness effect *>* 0.5 × 10^−3^). **Figure S7**- Adjustments to Δ*E* in a run of 1000 time steps for a population of 1000 individuals. **Figure S8**- Parameters conducive to adaptation in N=100 vs N=1,000 populations in the high mutation rate regime. **Figure S9**- Non-conducive DFEs. **Table 2**- Pearson’s correlation coefficients between DFE parameters (rows) and PC1, PC2 and PC3 N = 1000 populations. **Figure S10**- Three Principal Components explain DFE *conducivity*. **Figure S11**- Trade-off between the shape parameter and frequency and size of beneficial mutations for N = 1000 populations. **Figure S12**- Trade-off between the shape parameter and frequency and size of beneficial mutations for N = 100 populations in the high mutation rate regime. **Figure S13**- Time of minimum average fitness effects probability of retention of the locus in N=100 populations in the high mutation rate regime. **Figure S14**- Selection is essential for sustained fitness increase in the high mutation rate regime. **Table 3**- Pearson’s correlation coefficients between DFE parameters (rows) and time of minimum fitness for N = 100 and N = 1000 populations in the high mutation rate regime. **Figure S15**- Rare, large-effect positive jumps lead to sudden fitness increase in trajectories of adaptation. **Figure S16**- Scatter plot showing the distribution of minimum average fitness and the time of minimum average fitness for populations which eventually crossed the fitness threshold in the high mutation rate regime. **Figure S17**- Trajectory of population average fitness in the low mutation rate regime. **Figure S18**- Effect of locus deletion in N=1,000 populations in the high mutation rate regime. **Figure S19**- Histograms for correlations between expression level, adaptive value and fitness trajectories for populations where the fitness threshold of 0.1 was crossed. **Figure S20**- Histogram for correlations between random expression level (E rand) and fitness trajectories for populations where the fitness threshold of 0.1 was crossed. **Figure S21**- Scale of Δ*A* exponential fits as a function of the model parameter p.

## Notes

### Competing Interest Statement

The authors have declared no competing interest.

### Summary of Updates

Massive revision of the manuscript: title, abstract, results, analysis

