## Supplementary material for "Gene Birth in a Model of Non-genic Adaptation": SI

### Supplementary Information: Gene Birth in a model of Non-genic Adaptation

#### 1 Model time steps

**Table 1: Estimates of one model time-step in various species.** Mutation rates are in units of *per base pair per generation*. And the following references were used for generation times: *T. thermophila*: Cassidy-Hanley [2012], *S.cerevisiae, E.coli*: Milo et al. [2010], *S.pombe*: Petersen and Russell [2016], *C.reinhardtii*: Harris [2009], *P tetraurelia*: Ishida and Hori [2017] , *D.discoideum*: Fey et al. [2007], *S.typhimurium*: Silva et al. [2009], *D.melanogaster*: Fernández-Moreno et al. [2007], *A.thaliana*: Koornneef and Scheres [2001]. Most references give a range of values or multiple values of mutation rates and generation times depending on the culture conditions used. In such cases, we picked the average reported value at a single culture condition that is consistent between the studies reporting mutation rate and generation time in any one organism. The model time-step calculated in column 3 is the time it takes for 1 mutation to occur in a locus of 100 bp.

| Species | Generation time | mutation rate | reference (mutation rate) | **model time step** |
|---|---|---|---|---|
| *T.thermophila* | 2 hrs | $7.6 * 10^{-12}$ | Long et al. [2016] | $3 * 10^5$ yrs |
| *S.cerevisiae* | 90 min | $1.6 * 10^{-10}$ | Zhu et al. [2014] | $1.1 * 10^4$ yrs |
| *E.coli* | 20 min | $2 * 10^{-10}$ | Lee et al. [2012] | $1.9 * 10^3$ yrs |
| *S.pombe* | 2 hrs | $2 * 10^{-10}$ | Farlow et al. [2015] | $1.1 * 10^4$ yrs |
| *C.reinhardtii* | 6 hrs | $3 * 10^{-10}$ | Ness et al. [2012] | $2.3 * 10^4$ yrs |
| *P.tetraurelia* | 8 hrs | $2 * 10^{-11}$ | Sung et al. [2012] | $4.6 * 10^5$ yrs |
| *D.discoideum* | 4 hrs | $2.9 * 10^{-11}$ | Saxer et al. [2012] | $1.6 * 10^5$ yrs |
| *S.typhimurium* | 21 min | $3 * 10^{-9}$ | Lind and Andersson [2008] | 133 yrs |
| *D.melanogaster* | 10 days | $7 * 10^{-9}$ | Keightley et al. [2009] | $7.8 * 10^4$ yrs |
| *A.thaliana* | 6 weeks | $9 * 10^{-9}$ | Ossowski et al. [2010] | $2.4 * 10^4$ yrs |

#### 2 Probability of fixation is well approximated by relative fitness in the low mutation rate regime

In the low mutation rate regime, organisms undergo $\sim 10^8$ generations between two mutational time-steps. In these intervening generations, the variants of the locus compete among each other to leave progeny in the next generation. For most DFE parameter sets, a single variant is expected to be fixed in the population within these intervening generations; i.e., in the next model time-step only a single variant generated in the previous time-step is carried forward.

It is intractable to carry out simulations for $10^8$ generations between each of the 1000 model time-steps in a reasonable amount of time. Hence, we explored the dynamics of fixation within one model time-step in order to estimate the probability of fixation of any individual. We simulated the dynamics of selection in the intervening generations for all 432 parameter sets: For each parameter set, we generated $N = 1000$ mutations and simulated $10^8$ rounds of selection on these 1000 variants. We repeated these simulations 1000 times for each parameter set in order to obtain an estimate for the probability of fixation of any individual. We find that across parameters, a single variant gets fixed in the population within $1997.6 \pm 1078$ generations.

We wanted to estimate how the probability of fixation of any variant depends on its relative fitness in the population. As a simple first test, we checked whether the probability of fixation can be approximated by the relative fitness of the variants; i.e., we tested whether the distribution of individuals fixed in the detailed simulation described above matches that for when a single individual is randomly chosen according to its relative fitness.

We find that this is the case: for the 432 parameter sets, the KS-statistic for the distributions of relative fitness of individuals that get fixed in the detailed simulations versus the distribution of individuals simply picked according to their relative fitness is $0.038 \pm 0.011$ (p-values = $0.52 \pm 0.28$). The low values of the KS-statistics and high p-values indicate that the two sets of distributions are in reasonable agreement (FigS.1).

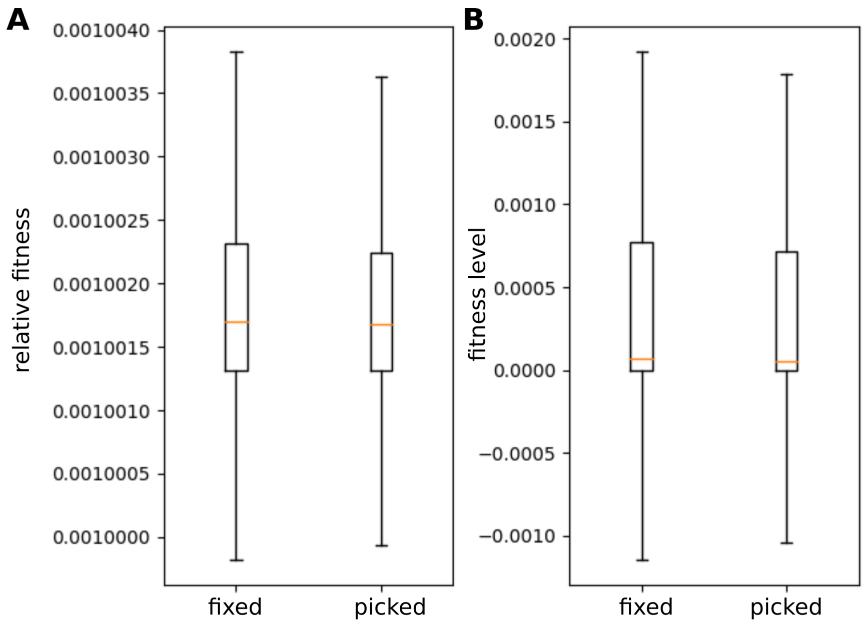


**FigS 1: The distribution of relative fitnesses and absolute fitness values found to fix in the detailed simulation (box label = 'fixed') is statistically similar to the distribution of relative fitnesses and absolute fitness values of individuals simply picked with a probability proportional to their relative fitness (box label = picked)**. This figures shows results for the *Chlamydomonas* parameters (p=0.001, n=-0.01, f=0.75, s=0.3, d=0). The boxes represent the distributions, orange lines represent medians, and whiskers represent quartiles. (A) Distributions of relative fitness. (B) Distributions of absolute fitness values.

### 3 The effect of genomic background on the adaptation of single loci

In the following simulations, we test how the evolutionary outcomes can change if we consider the simultaneous evolution of multiple genomic loci. We perform these simulations for the high mutation rate regime.

#### 3.1 Multi-locus model

In the model, we make the simplifying assumption of the fitness effects of the genetic background being constant over the time-scales being considered. This assumption requires that the evolution of the rest of the genome is very fast compared to that of the locus of interest. This is plausible for organisms with very large genomes, such that the number of mutations incident on the rest of the genome is many times over the number of mutations incident on the locus of interest.

We test this assumption in a more comprehensive multi-locus model, where we consider $N_l = 200$ loci, each of which is randomly assigned a characteristic distribution of fitness effects (DFE) of mutations. These loci were randomly assigned DFEs with parameters picked from $p = 0.001, 0.003, 0.005$, $n = 0.001, 0.005, 0.01$, $f = 0.25, 0.5, 0.75$, $s = 0.3, 0.6, 0.9$. Multiple loci in this *genome* have the same DFE parameters, thereby allowing us to compare their outcomes (FigS.2(A)). Particularly, 4 out of the 200 loci had the *Chlamydomonas* DFE parameters. At each time step, a randomly chosen locus incurs a mutation in each individual of the population. Fitness of an individual is the sum of fitness contributions from all loci. Similar to the main model, the number of offspring an individual leaves is proportional to the relative fitness. Individuals with even a single locus that has a fitness contribution less than -1 are inviable.

For this simulation, we used a population size of $N = 1000$, and ran for $T = 100,000$ time steps. In this model, the number of mutations incident on any one locus across the population during the simulation is on average $(N \times T)/N_l$. This is half the number of mutations encountered by the locus in the main model. Therefore, in order to compare the ability of loci to adapt in this multi-locus model with that in the main model, we count the number of loci with a given DFE that cross the fitness threshold of 0.05.

The DFE parameters for loci that crossed the 0.05 threshold in this simulation almost exactly match the DFE parameters of loci that cross the 0.1 threshold in the main model (FigS.2(B)). In particular, all 4 loci with the *Chlamydomonas* DFE parameters cross the threshold in the multi-locus simulation. This indicates that our treatment of the genomic background is reasonable for the purpose of this study.

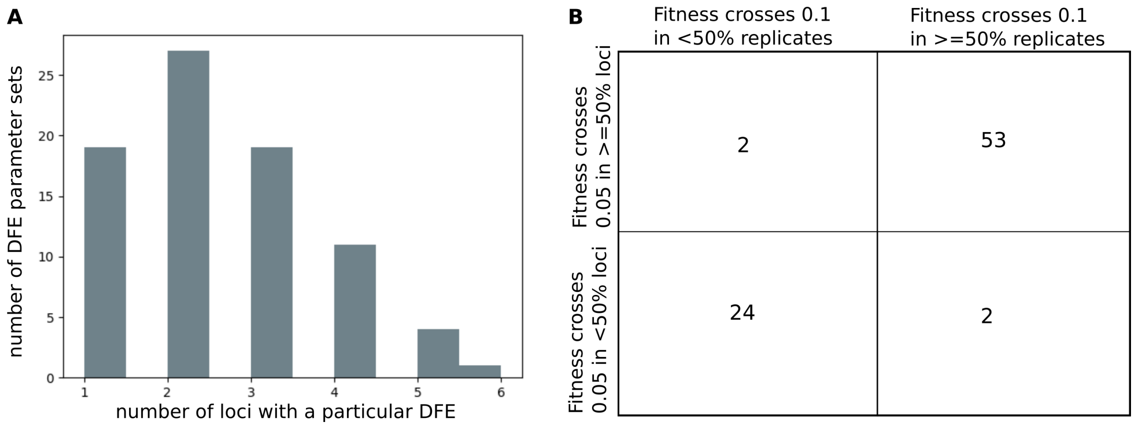


**FigS 2: Comparison of adaptation of loci in the main model versus the multi-locus model.** (A) Histogram for the number of loci out of 200 that have a particular set of DFE parameters $(p, n, f, s)$. There is at least one locus corresponding to each of the 81 DFE parameter sets tested in this study. (B) Comparison of the DFE parameter sets for which fitness thresholds were crossed in the main model versus the multi-locus model.

##### 3.2 Evolution of non-genic locus in the presence of an established gene

In real organisms, *de novo* gene birth occurs in the presence of well-established genes. While the multi-locus model of the previous section indicates that the presence of other *de novo* genes evolving independently in the same genome should not skew the outcomes for *de novo* gene evolution at any single locus, we wanted to test how the presence of an evolving established gene might change these outcomes.

Here, we consider two loci, one of which is initially non-genic ($F_d^{ini}enovo = 0$) and the other is an established gene ($F_e^{ini}st = 0.5$). We use a fixed conservative DFE to draw the effect of mutations on the established gene ($p = 0.001, n = 0.001, f = 0.25, s = 0.1$), and we survey all 108 DFEs used in the main study for the non-genic locus. In these simulations, population size $N = 1000$, and we simulate for $T = 1000$ time steps. At each time step, a randomly chosen locus (non-genic or established gene) incurs a mutation in each individual of the population. Fitness of an individual is the sum of fitness contributions from both loci. Similar to the main model, the number of offspring an individual leaves is proportional to the relative fitness. Individuals with even a single locus that has a fitness contribution less than -1 are inviable. For each of the 108 non-genic DFE parameter sets, we simulate 25 replicate populations. We count the number of replicates for which the initially non-genic locus crosses the fitness threshold of 0.1 at the end 1000 time steps. We find that The DFE parameters for loci that crossed the 0.1 threshold in this simulation almost exactly match the DFE parameters of loci that cross the 0.1 threshold in the main model(FigS.3(B)).

These results indicate that independently evolving loci, irrespective of initial conditions and DFE parameters should not change the outcomes of evolution for any single locus. It is therefore likely that only loci which interact ,for example through genetic linkage or epistasis, can influence evolutionary outcomes.

##### 3.3 Diminishing returns epistasis can reduce ability of loci to adapt

In the model, the DFE of the intergenic loci of interest remain constant over time. However, over the evolutionary timescales that the model captures, the DFE of a locus is likely to vary. An important manner in which DFE of established genes vary over time is through diminishing returns epistasis, where the strength/number of beneficial mutations decreases as the fitness contribution of genes increases. Implementing diminishing returns epistasis in our model is made difficult due to important unknown quantities, especially for non-genic loci. These unknown quantities include (1) the functional form of diminishing returns (is it the effect size, or the number of beneficial mutations that decreases with fitness? Do the positive effects decrease linearly, or non-linearly?), and (2) how sharply do positive effect mutations diminish with increasing fitness?

A detailed exploration of the relevant forms of diminishing returns epistasis which impact *de novo* gene birth will be a very important additional layer to consider. However, due to the lack of empirical studies on non-genic loci, this exploration is beyond the scope of the current work. For these reasons, we proceed in our model with the assumption

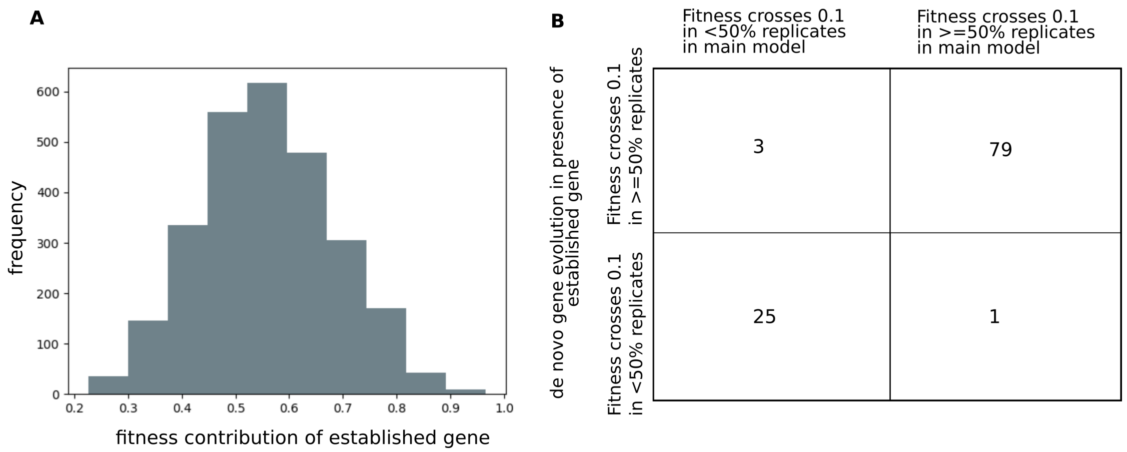


**FigS 3: Comparison of adaptation of loci in the main model versus a model where *de novo* gene evolution occurs in the presence of an independently evolving established gene.** (A) Histogram of final fitness contribution of established gene at the end of the simulation. (B) Comparison of the DFE parameter sets for which fitness thresholds were crossed in the main model versus the model with established gene.

that at early stages of gene birth, which this model captures, the fitness contribution of the emerging gene is very low and is not subject to diminishing returns epistasis.

Below we demonstrate how diminishing returns epistasis can reduce the frequency of gene birth in the model in the high mutation rate regime. In these simulations, the parameter p (mean effect size of beneficial mutations) reduces as F (fitness of the individual) increases. We assume that p remains constant whenever F is very low (F < 0.01), p reaches a constant and very small value of 0.0005 (which is effectively neutral) when F crosses a threshold $F_{high}$, and p decreases linearly for $0.01 < F < F_{high}$. We simulate systems with initial DFE (p=0.001, f=0.75, n=0.01, s=0.3) and test two values of $F_{high}$: 0.25 (FigS.4(A)), and 0.5 (FigS.4(B)).

We find that fitness trajectories in these simulations are very noisy, since mutations are now drawn from different DFEs for each individual in the population. As expected, p decreases slower for $F_{high} = 0.5$, and consequently populations reach higher values of average fitness.

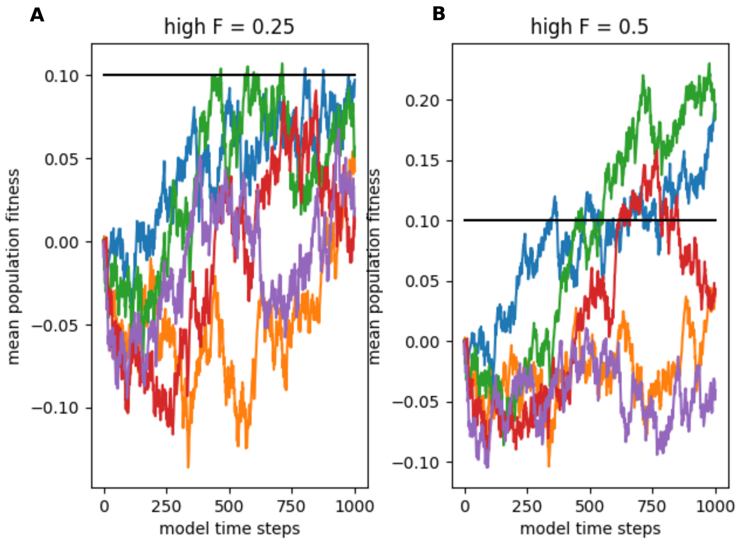


**FigS 4: Trajectories of population averaged fitness subject to diminishing returns epistasis.** Colors indicate trajectories for 5 independent simulations with the same initial DFE parameters: (p=0.001, n=0.01, f=0.75, s=0.3). The horizontal black line indicates fitness = 0.1. (A) simulations with $F_{high} = 0.25$. (B) simulations with $F_{high} = 0.5$.

#### 4 A majority of mutations sampled in this study are neutral

Most mutations incident on non-genic regions are expected to be nearly neutral. Therefore, a good set of DFE parameters to test in this model should include those where a majority of the mutations are neutral. Neutral mutations are those that are not visible to natural selection. In infinite populations, all mutations, no matter how small their effect, do change the course of evolution. But in finite populations, which are subject to random drift, small mutations are effectively neutral. Here, we test the magnitude of mutational fitness effect which is effectively neutral in a N=1000 population.
In order to do this, we simulate the evolution of a N = 1000 population, where initially, 900 individuals are wild-type (F = 1) and 100 individuals are mutants ($F = 1 + mut$). We want to find the value of $mut$ for which the course of evolution of populations with and without selection are statistically indistinguishable. We carry out two parallel simulations for the population: (A) with selection (high mutation rate regime): At each time-step, the probability that an individual i with fitness $F_i$ leaves an offspring is proportional to $F_i$, and (B) no selection: At each time-step, the probability that an individual i with fitness $F_i$ leaves an offspring is 0.5. We carry out these parallel simulations for 1000 time-steps, and count the number of mutants in the population in the end.
We repeat these parallel simulations for 1000 replicate populations. We find in this analysis that mutations with a fitness effect of $0.5 \times 10^{-3}$ are effectively neutral for N = 1000 populations (FigS.5).

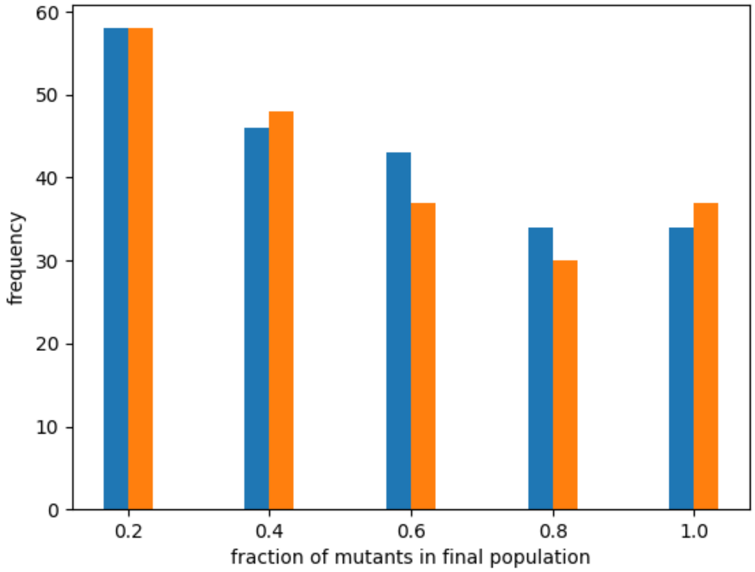


**FigS 5: Neutrality of mutants with fitness effect** $= 0.5 \times 10^{-3}$ **in N=1000 populations.** The x-axis represents the fraction of mutants in the final population after 1000 time-steps ($mutant - fraction$). The x-axis is binned, and we count populations with $x < mutant - fraction <= x + 0.2$ in the same bin. The y-axis represents the number of replicate populations out of 1000 which end up with the same $mutant - fraction$. Blue bars indicate results of simulations with selection and orange bars indicate results of simulations with no selection. When the effect of the mutation $mut = 0.5 \times 10^{-3}$, the results of the two simulations are indistinguishable (two sample KS-test: test statistic = 0.009, p-value = 0.99). The high p-value indicates that it is very likely that the two samples are drawn from the same distribution.

We then wanted to count the number of neutral, positive effect and negative effect mutations which the DFEs tested in our work produce. To do this, we draw $10^6$ independent mutations from each of the DFE parameters used in this study. We then count the number of neutral mutations ($-0.5 \times 10^{-3} < mut \leq 0.5 \times 10^{-3}$), positive effect mutations ($mut > 0.5 \times 10^{-3}$), and negative effect mutations ($mut \leq -0.5 \times 10^{-3}$). We find that mutations drawn using DFE parameters tested in the model, the proportion of positive effect mutations can be much lower than what is suggested by the parameter $f$ (fraction of beneficial mutations) (FigS.6(A)). We also find that many DFE parameters which produced very small proportions of positive effect mutations show adaptation in the high mutation rate regime (FigS.6(B)).

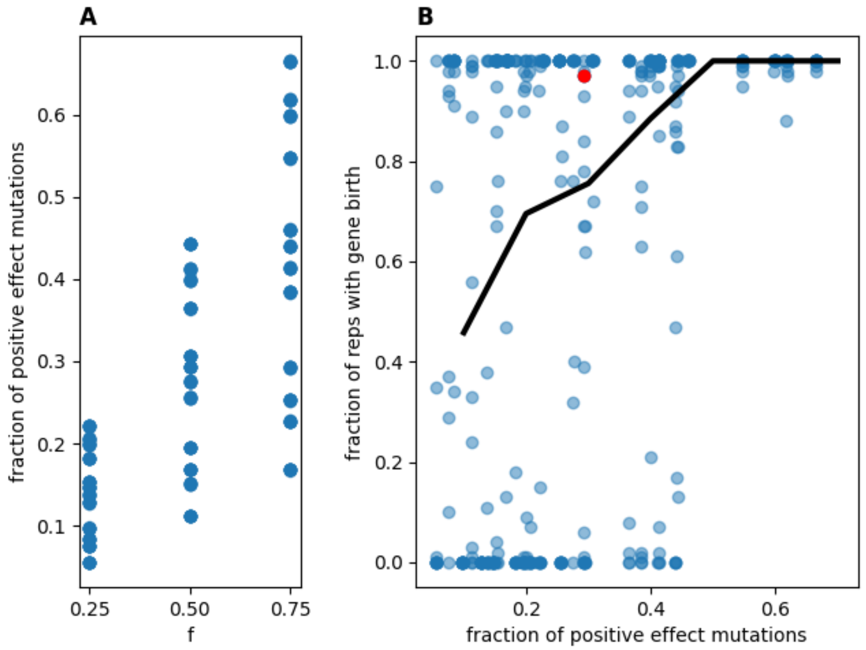


**FigS 6: Fraction of positive effect mutations (fitness effect $> 0.5 \times 10^{-3}$).**(A) Fraction of positive effect mutations out of $10^6$ mutations drawn using DFEs used in our work. The x-axis represents values of the model parameter f (fraction of beneficial mutations) and the y-axis represents $f_p$, the fraction of non-neutral, positive effect mutations (fitness contribution >0.0005). (B) Fractions of replicate populations with gene birth in the high mutation rate regime versus fraction of positive effect mutations. Each point represents a distinct DFE parameter set $(p, n, s, f)$. The x-axis represents $f_p$ for all DFE parameter sets used in our work, and the y-axis represents the fraction of replicate populations with gene birth for the corresponding DFE parameter set (deletion probability, $d = 0$ for all points shown). The black line indicates average fraction of replicates with gene birth for all DFE parameter sets with values of $f_p$ given by the x-axis. The red point indicates values of $f_p$ and fractions of replicates with gene birth in the high mutation rate regime for the parameter set (p=0.001, n=0.01, f=0.75, s=0.3), which corresponds to the DFE closest to that of *Chlamydomonas reinhardtii* reported in (Böndel et al. [2019])

### 5 Deviation of $\Delta E$ distribution from the intended power law

The distribution of $\Delta E$ reported in Vaishnav et al. [2022] provides us with estimates of the magnitudes of changes in expression level due to mutations. Such changes in expression level are highly likely to be correlated with corresponding changes in fitness levels due to the mutation. In our model, due to unavailability of data on these correlations, we draw $\Delta E$ and $\Delta f$ independently of each other.

Importantly, we assume that $\Delta E$ is equally likely to be positive or negative. In our simulations, we initialize the locus with a very low value of expression level (=0.001), and expression level is not allowed to be negative. Therefore, we make adjustments to the value of $\Delta E$ if the expression level drops too low (see main text: Method to update expression level and adaptive value). Such adjustments necessarily lead to small deviations of the $\Delta E$ distribution from the power law distribution (FigS.7).

### 6 Adaptive evolution of non-functional loci as a function of model parameters and population size

#### 6.1 Adaptive evolution of non-functional loci at population size N=100

We also performed simulations for N=100 populations in the **high mutation rate regime**. Our results remain qualitatively the same: the probability of the fitness contribution of the locus crossing the threshold of 0.1 is positively correlated with $p$ and $f$, and negatively correlated with $n$ and $s$ (FigS.8(A), inset). And similar to the N=1000 populations, time of minimum fitness is correlated with the model parameters (Table.S3).

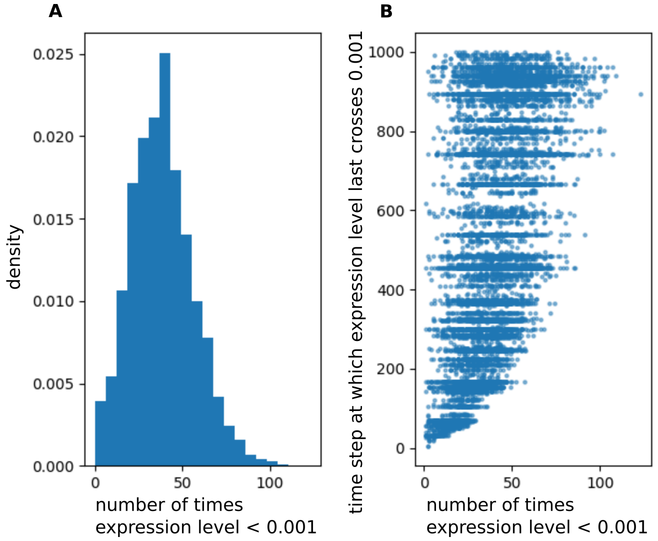


**FigS 7: Adjustments to $\Delta E$ in a run of 1000 time steps for a population of 1000 individuals.** Each individual is treated independently of the others. (A) Histogram for the number of times adjustments are applied to $\Delta E$. (B) Scatter plot for number of times an adjustment was applied (x-axis) versus the final time step at which the adjustment was applied.

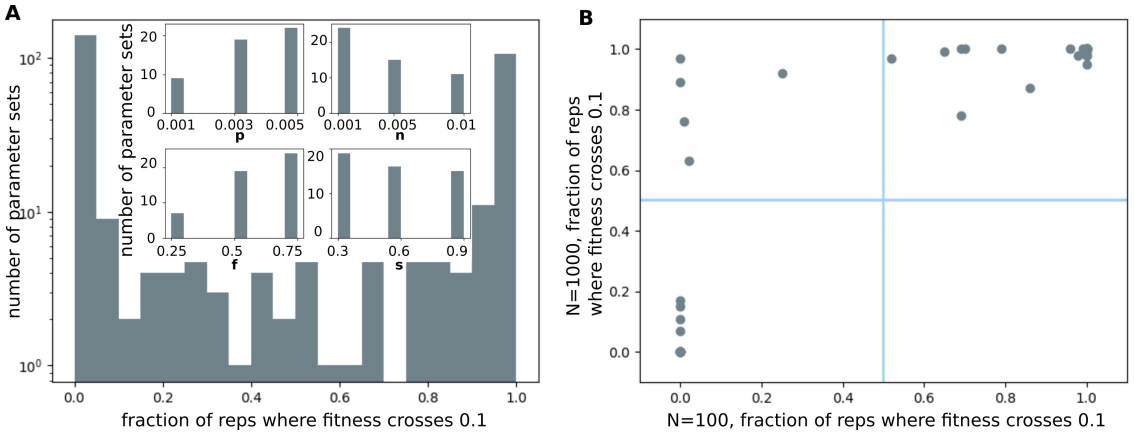


**FigS 8: Parameters conducive to adaptation in N=100 vs N=1000 populations in the high mutation rate regime.** (A)Histogram for the fraction of replicate populations that cross the fitness threshold for various parameter sets $(p, n, f, s)$, for $d = 0$ and population size $N = 100$. Inset: Histograms for the number of parameter sets with given values of parameters $p, n, f$, or $s$ for which more than half of the replicate populations cross the fitness threshold. (B) Scatter plot comparing the fraction of replicate populations that cross the fitness threshold for various parameter sets for N=100 (x-axis) and N=1000 (y-axis). Each point represents a parameter set. For points in the upper right quadrant indicate parameter sets where $> 50\%$ of replicates cross the threshold in both N=100 and N=1000 populations. Points in the upper left quadrant cross the threshold only in N=1000 populations.

##### 6.2 Non-conducive DFE parameters

In the high mutation rate regime, $28/108$ of the sampled DFE parameters were not *conducive* to gene birth. As can be seen from the histogram in Fig.3(A), less than 0.2 of replicate simulations with these DFE parameters resulted in the fixation of a variant of the locus that had $F >= 0.1$. All but one of these DFE parameters represent very conservative scenarios that produce more numerous negative than positive effect mutations. In FigS.9 (blue bars), we show how DFEs with low values of parameters $p, f$ and high values of $n, s$ tend to be *non-conducive* in the high mutation rate regime.

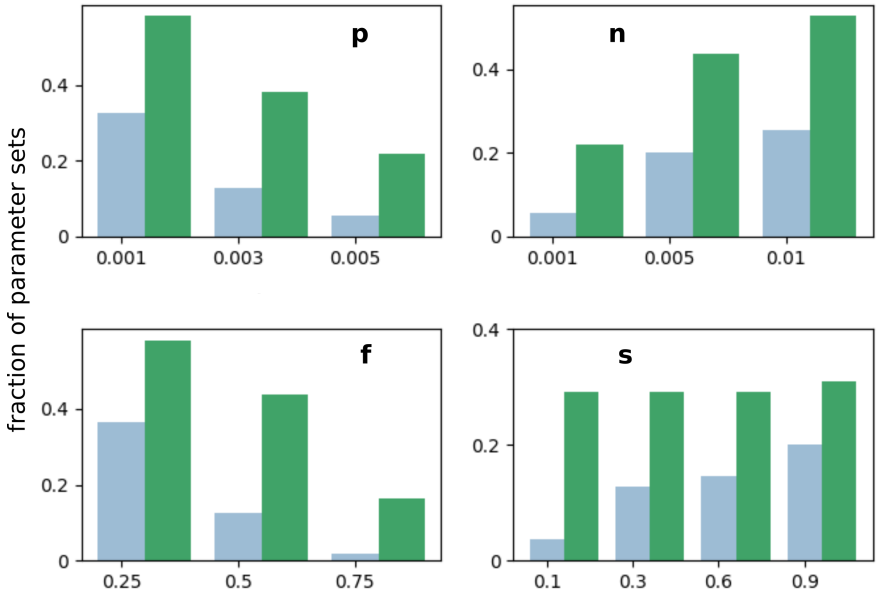


**FigS 9: Non-conducive DFEs.** Histograms for the fraction of parameter sets with given values of parameters p, n, f , or s for which less than half of the replicate populations cross the fitness threshold. Blue bars: high mutation rate regime; green bars: low mutation rate regime.

In contrast, in the low mutation rate regime, DFEs with low values of parameters $p$, $f$ and high values of $n$ tend to be *non-conducive*, but there is no effect of the shape parameter $s$ (FigS.9, green bars).

##### 6.3 Interactions between model parameters

Often, measurable parameters that are used to characterize biological systems can be correlated with each other such that the actual dimensionality of the system is much lower than the number of these measured parameters (Eckmann and Tlusty [2021]).
In order to test how DFE parameters interact among each other, we performed Principle Components Analysis over 8 variables: the proportions of positive effect and negative effect mutations for each DFE; the average magnitudes of positive effect and negative effect mutations; the largest magnitudes of positive effect and negative effect mutations; and the smallest magnitudes of positive effect and negative effect mutations produced by each DFE. From the elbow plot (FigS.10(A)), we see that individually, the first 3 principle components (PCs) explain large fractions of the variance, and taken together they explain 77.5% of the variance. The first 5 PCs together explain 94% of the variance.

In the high mutation rate regime, DFE *conducivity* (the fraction of replicate populations in which a variant of the locus with fitness >= 0.1 gets fixed) is significantly correlated with PC1, PC2 and PC3 (FigS.10(B)). In contrast, DFE *conducivity* in the low mutation rate regime correlates significantly with PC2 and PC3 but not PC1 (FigS.10(C)). These PCs are highly correlated with all model parameters, particularly, PC1 is significantly correlated with the shape parameter, s (TableS.2).

**Table 2: Pearson's correlation coefficients between DFE parameters (rows) and PC1, PC2 and PC3 for N = 1000 populations.** The numbers in the parentheses indicate p-values.

| DFE parameters | Correlation coeff. (PC1) | Correlation coeff. (PC2) | Correlation coeff.(PC3) |
|---|---|---|---|
| $p$ | 0.29 (0.002) | 0.16 (0.085) | 0.41 ($1.1 \times 10^{-5}$) |
| $n$ | 0.30 (0.001) | -0.14 (0.16) | -0.75 ($5.4 \times 10^{-21}$) |
| $f$ | -0.07 (0.49) | -0.86 ($5.3 \times 10^{-32}$) | -0.19 (0.05) |
| $s$ | -0.76 ($2.9 \times 10^{-21}$) | -0.02 (0.76) | -0.13 (0.18) |
| *conducivity* (Hi-mut) | 0.25 (0.01) | -0.50 ($3.6 \times 10^{-8}$) | 0.49 ($6.6 \times 10^{-8}$) |
| *conducivity* (Lo-mut) | 0.01 (0.90) | -0.47 ($3.7 \times 10^{-7}$) | 0.67 ($1.5 \times 10^{-15}$) |

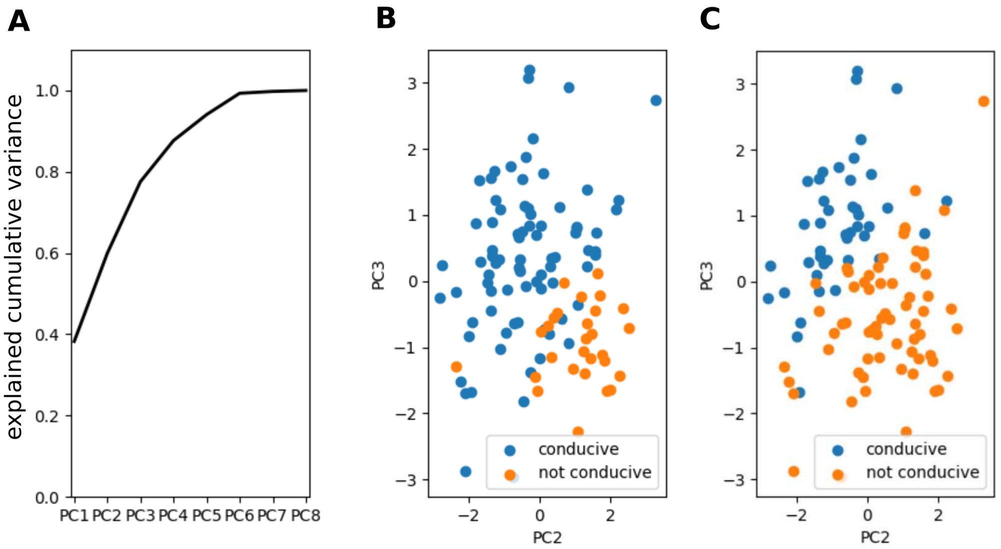


**FigS 10: Three Principle Components explain DFE *conducivity***(A) PCA elbow plot indicates that principle components 1,2 and 3 explain 77.5% variance. (B,C) Scatter plot of values of PC2 and PC3 for each of the 108 DFE sampled in this study. Blue points indicate DFEs with $conducivity \geq 0.5$, and orange points indicate DFEs with $conducivity < 0.5$ for (B) high mutation rate regime, (C) low mutation rate regime.

This dependence of *conducivity* on the shape parameter in the high mutation rate regime allows DFEs that produce small and infrequent beneficial mutations (small p,f) to still lead to locus adaptation if the magnitude of the shape parameter (s) is also low (FigS.11, FigS.12).

##### 6.4 Differences in patterns of evolution due to population size

There are substantial differences in the fitness trajectories of N=100 populations due to increased effect of random drift (FigS.13(A)). While for most DFEs, spontaneous mutations are mostly deleterious, and selection is required to cross the fitness threshold of 0.1, the level of drift in N=100 populations is able to overcome this effect (FigS.14). On the flip side, drift also leads to the populations being more susceptible to locus deletion: While in the absence of locus deletion, the fitness threshold of 0.1 was crossed in more than 50% of replicate N=100 populations for 50 out of 81 parameter sets, including the *Chlamydomonas* parameters, in the presence of locus deletion, the number of parameter sets for which the locus survives deletion is much lower (FigS.13(B)).

#### 7 A model of DFE with extremely rare positive jumps yields noisy trajectories

We tested a different mechanism of *de novo* gene birth in the high mutation rate regime: In these simulations, we use a DFE such that most mutations are neutral ($\sim$70% of the mutations are neutral for the chosen DFE parameters), and there are very few positive effect mutations ($\sim$5%). We verify that, on their own, mutations drawn from this DFE do not lead to gene birth (Fig.15(A)).

We then augmented this DFE with rare (probability=0.0001), large and positive jumps of fixed effect size (0.15). These rare positive jumps represent molecular mechanisms that could potentially be involved in *de novo* gene birth but were not considered in our main model. We find that the trajectories of average population fitness with this new mechanism are noisier than the standard case in our study, and can be punctuated by noticeable jumps (Fig.15(B): red and orange trajectories). At the same time, we note that, although noisier, the general tendency of the trajectories is similar to the model considered in the main text. Hence, presence of rare, large positive jumps is sufficient for adaptation, and adaptation can also result from mixtures of different molecular mechanisms.

#### 8 Time of minimum fitness is correlated with model parameters in the high mutation rate regime

In the high mutation rate regime, the dynamics of population averaged fitness switches from a mutation dominated phase to a selection dominated phase at the *time of minimum fitness*. The time of minimum fitness and the minimum

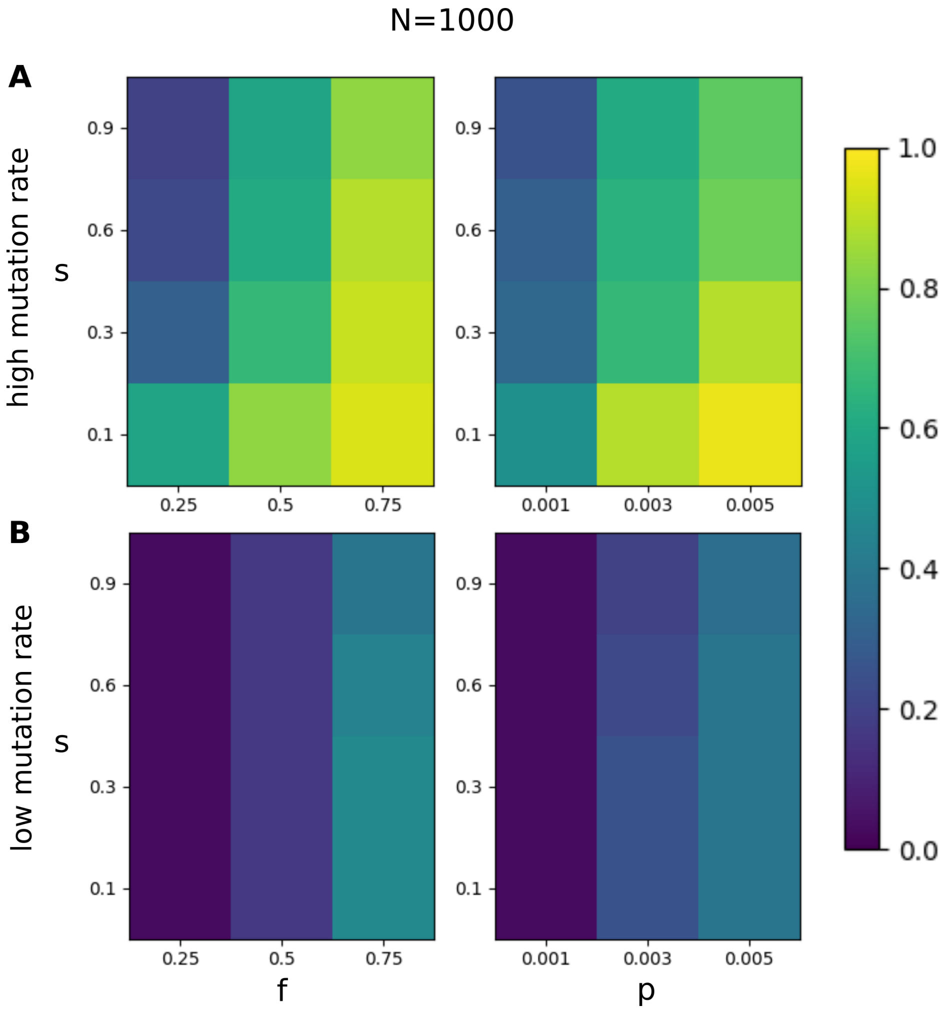


**FigS 11: Trade-off between the shape parameter and size of beneficial mutations for N = 1000 populations.** In the heatmaps, rows indicate values of the shape parameter, columns indicate values of (right panels) mean size of beneficial mutations $p$, (left panels) fraction of beneficial mutations *f*. Colors indicate the fractions of systems such that at least 50% of the replicate populations crossed the fitness threshold of 0.1. (A) Heatmaps for the high mutation rate regime. (B) Heatmaps for the low mutation rate regime.

average fitness in the population are strongly correlated (Pearson's correlation coefficient = -0.87, p-value = $5.5 \times 10^{-34}$. See FigS.16). The *time of minimum fitness* is also correlated with model parameters (TabS.3).

#### 9 Fitness trajectories in the low mutation rate regime do not show consistent adaptation

In the low mutation rate regime, there is no 'selection dominated phase'. Even for *conducive* DFE parameters, fitness trajectories do not show a phase of persistent fitness increase (FigS.17).

#### 10 ODE model describing the effect of locus deletion in the high mutation rate regime

We describe here the effect of locus deletion on the evolutionary dynamics of non-functional loci in the high mutation rate regime in terms of competition between individuals that contain the locus, and those that have lost the locus. In the reduced ODE model described below, $x$ is the fraction of the population that contains the locus, and $y$ is the fraction that has lost the locus. Initially, $x = 1$ and $y = 0$. An individual containing the locus can lose it with probability $d$. For the fraction of the population that contains the locus, locus fitness $f$ follows a parabolic trajectory with time $t$:

$df/dt = a * (2t - b)$

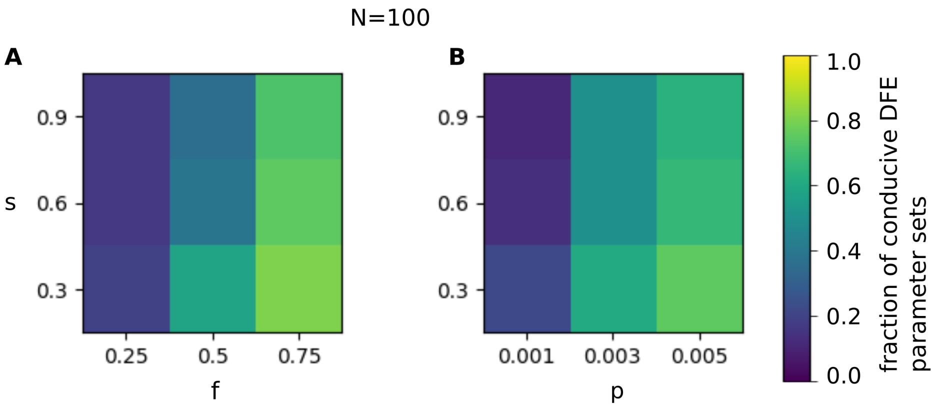


**FigS 12: Trade-off between the shape parameter and frequency and size of beneficial mutations for N = 100 populations in the high mutation rate regime.** In the heatmaps, rows indicate values of the shape parameter, columns indicate values of (A) fraction of beneficial mutation $f$, (B) mean size of beneficial mutations $p$. Colors indicate the fractions of systems such that at least 50% of the replicate populations crossed the fitness threshold of 0.1.

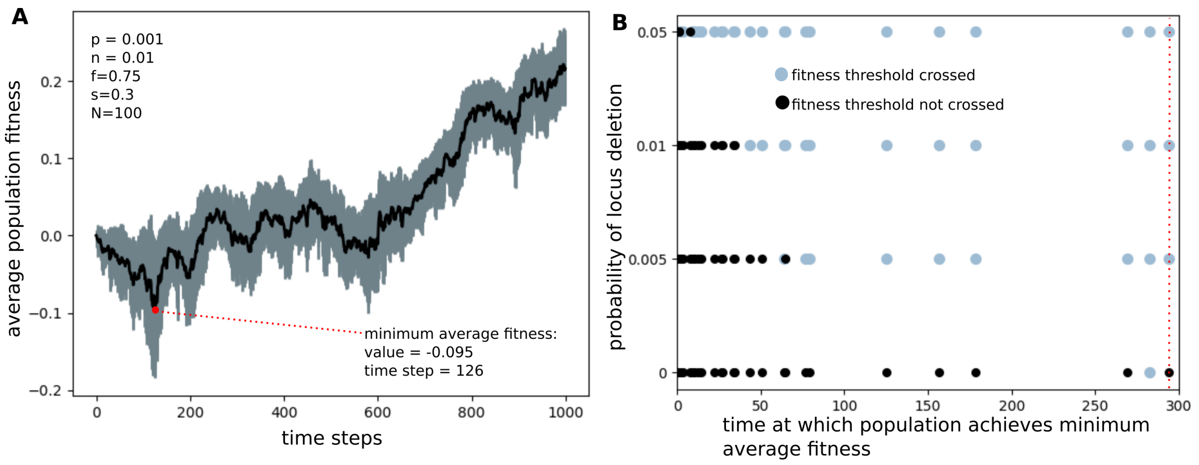


**FigS 13: Time of minimum average fitness effects probability of retention of the locus in N=100 populations in the high mutation rate regime.**(A) The trajectory of population average fitness in one of the replicate populations with *Chlamydomonas* DFE parameters, with no locus deletion. Average fitness is indicated by the black line, and the grey shading represents standard deviation. The red point indicates the time step at which average fitness has the minimum value. Parameters used to generate the fitness trajectory are given in the figure. Here, the locus deletion probability $d = 0$. (B) Scatter plot showing how populations where minimum average fitness is achieved later are more likely to be affected by locus deletion. Each point represents a parameter set. The x-axis indicates the time at which minimum average fitness is achieved averaged over all the replicate populations with the same DFE parameters $(p, n, f, s)$. Black points represent parameter sets where $\geq 50\%$ of the replicate populations cross the fitness threshold, and blue points represent parameter sets where $< 50\%$ of the replicate populations cross the fitness threshold. The dotted red line indicates the *time of minimum average fitness* for *Chlamydomonas* parameters.

This trajectory resembles the model's average fitness trajectories of populations which cross the fitness threshold (see main text: Fig.2(B)) in that they are characterized by two parameters $a$ and $b$ which govern the *minimum fitness* ( $= -(a * b^2)/2$) and *time of minimum fitness* ($= b/2$) respectively (FigS.18(A)).

Competition between individuals is captured by the following equations that govern the growth of the fractions of the populations $x$ and $y$:

$dx/dt = x * (1 + f) * (1 - x^2) - d * x$ — (1)

$dy/dt = y * (1 - y^2) - f * x + d * x$ — (2)

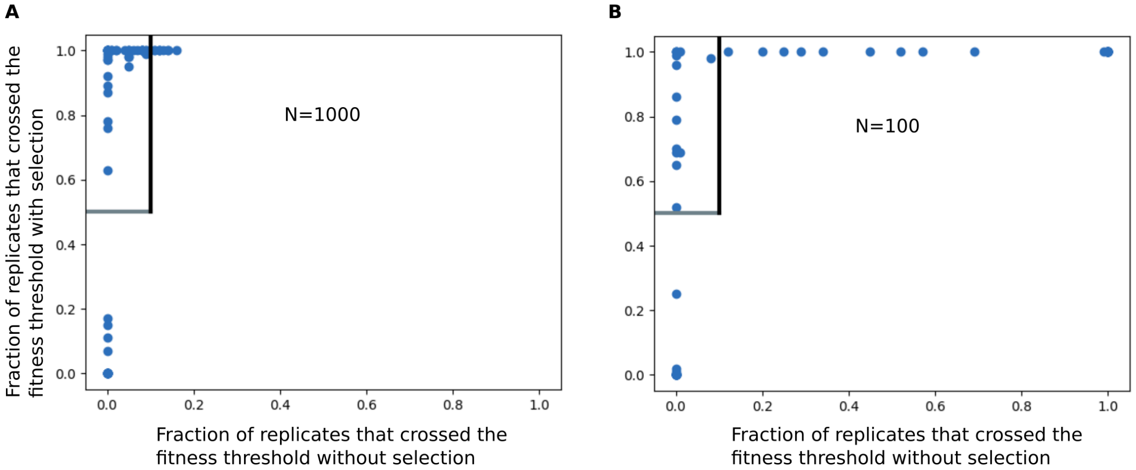


**FigS 14: Selection is essential for sustained fitness increase in the high mutation rate regime.** Scatter plots for the fraction of replicate populations that cross the fitness threshold of 0.1 with (y-axis) versus without (x-axis) selection. Each point represents a distinct parameter set. (A)N=1000, (B) N=100.

**Table 3: Pearson's correlation coefficients between DFE parameters (rows) and time of minimum fitness for N = 100 and N = 1000 populations in the high mutation rate regime.** The numbers in the parentheses indicate p-values.

| | N=100 | N=1000 |
|---|---|---|
| $p$ | -0.44 ($4.5 \times 10^{-5}$) | -0.41 ($9.3 \times 10^{-6}$) |
| $n$ | 0.29 (0.01) | 0.26 ($5.8 \times 10^{-3}$) |
| $f$ | -0.56 ($5 \times 10^{-8}$) | -0.51 ($1.9 \times 10^{-8}$) |
| $s$ | 0.076 (0.5) | 0.21 (0.03) |

In the right hand sides equations (1) and (2), the first terms capture logistic growth with adjustments to keep values of $x$ and $y$ between 0 and 1. Fitness of the locus, $f$, introduces competition: positive $f$ favors the growth of $x$ and hinders $y$. The last terms represent the effect of locus deletion; individuals are transferred from population $x$ to population $y$ at a rate $d$.

We run the ODE for a time $t = 1000$. We then determine whether the locus would survive in a population of size $N$ by asking if at least one individual with the locus survives in the population, i.e. $x * N >= 1$. We find that for populations which achieve minimum fitness at later times, and have a lower value of minimum fitness, the locus only survives for small values of $d$ (FigS.18(B)). Since *time of minimum fitness* and *minimum fitness* are correlated in the model, this explains how locus deletion blocks the adaptive evolution of non-functional loci.

#### 11 Expression level and adaptive value as drivers of fitness trajectories in the high mutation rate regime

In our simulations, for most populations which cross the fitness threshold of 0.1, population average trajectories of both expression level trajectories and adaptive value trajectories were positively correlated with the average fitness trajectory. Nevertheless, a comparison of correlation coefficients across all populations suggests that fitness change is more strongly driven by adaptive value (FigS.19).

#### 12 Distributions of $\Delta A$ in the high mutation rate regime

The exponential distribution is a good fit for $\Delta A$ values across all parameter sets. The scale of the best fit distribution varies according to the parameter values; for example, the scale increases on average as a function of the size of beneficial mutations $p$ (FigS.21).

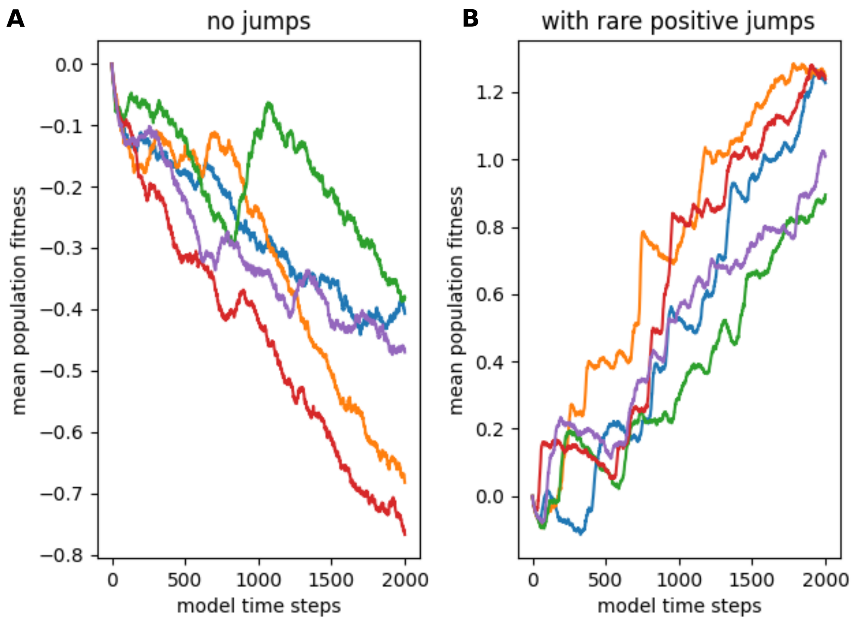


**FigS 15: Rare, large-effect positive jumps lead to sudden fitness increase in trajectories of adaptation in the high mutation rate regime.** Trajectories of population averaged fitness using a DFE with parameters (p=0.001, n=0.005, f=0.25, s=0.1). Colors indicate trajectories for 5 independent simulations with the same DFE parameters. The plot on the left represents trajectories where only small mutations drawn from the given DFE occur. The plot on the right represents a model where in addition to small mutations, rare and large positive jumps also occur.

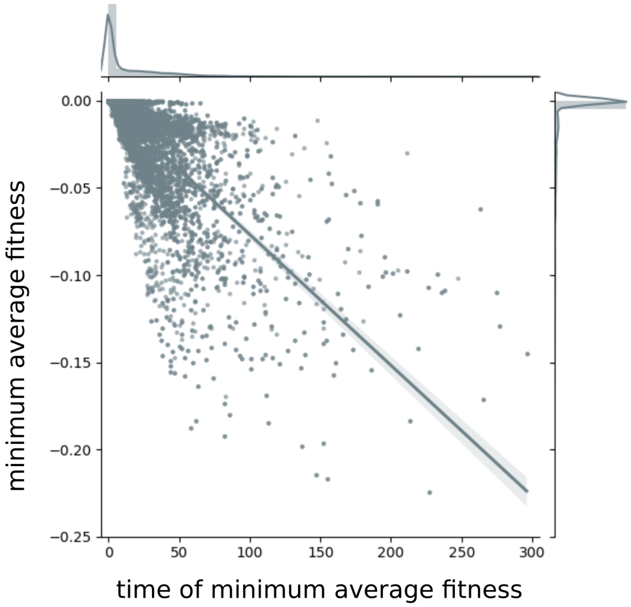


**FigS 16: Scatter plot showing the distribution of minimum average fitness and the time of minimum average fitness for populations which eventually crossed the fitness threshold in the high mutation rate regime.** The histograms are the marginal distributions of time of minimum fitness and minimum fitness.

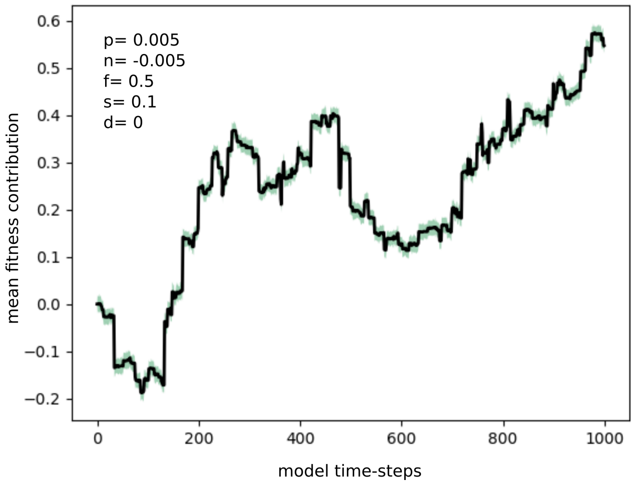


**FigS 17: Trajectory of population average fitness in the low mutation rate regime** in one of the replicate populations with DFE parameters as indicated in the legend, and no locus deletion ($d = 0$). This parameter set corresponds to the most stringent DFE for which the population crossed the fitness threshold. Average fitness is indicated by the black line, and shading represents standard deviation.

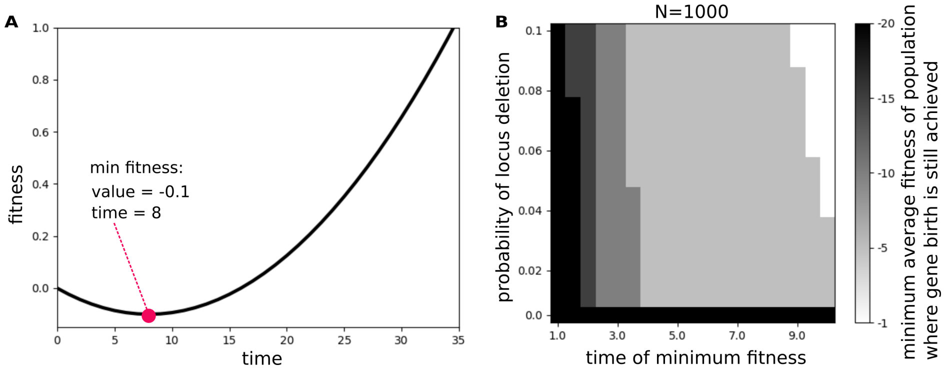


**FigS 18: Effect of locus deletion in N=1000 populations in the high mutation rate regime.** (A) Fitness trajectory in the ODE model for parameter values $a = 0.1$ and $b = 2$.(B) Heatmap of survival of the locus in N=1000 populations. The rows represent the probability of locus deletion ($d$) and the columns represent the time of minimum fitness. Colours indicate the most negative minimum fitness which survives in the population.

Ron Milo, Paul Jorgensen, Uri Moran, Griffin Weber, and Michael Springer. Bionumbers—the database of key numbers in molecular and cell biology. *Nucleic acids research*, 38(suppl_1):D750–D753, 2010.

Janni Petersen and Paul Russell. Growth and the environment of schizosaccharomyces pombe. *Cold Spring Harbor Protocols*, 2016(3):pdb–top079764, 2016.

Elizabeth H Harris. *The Chlamydomonas Sourcebook: Introduction to Chlamydomonas and Its Laboratory Use: Volume 1*, volume 1. Academic press, 2009.

Masaki Ishida and Manabu Hori. Improved isolation method to establish axenic strains of paramecium. *Japanese Journal of Protozoology*, 50(1-2):1–14, 2017.

Petra Fey, Anthony S Kowal, Pascale Gaudet, Karen E Pilcher, and Rex L Chisholm. Protocols for growth and development of dictyostelium discoideum. *Nature protocols*, 2(6):1307–1316, 2007.

Roberta Ribeiro Silva, Célia Alencar Moraes, Josefina Bessan, and Maria Cristina Dantas Vanetti. Validation of a predictive model describing growth of salmonella in enteral feeds. *Brazilian Journal of Microbiology*, 40(1):149–154, 2009.

Miguel Angel Fernández-Moreno, Carol L Farr, Laurie S Kaguni, and Rafael Garesse. Drosophila melanogaster as a model system to study mitochondrial biology. In *Mitochondria*, pages 33–49. Springer, 2007.

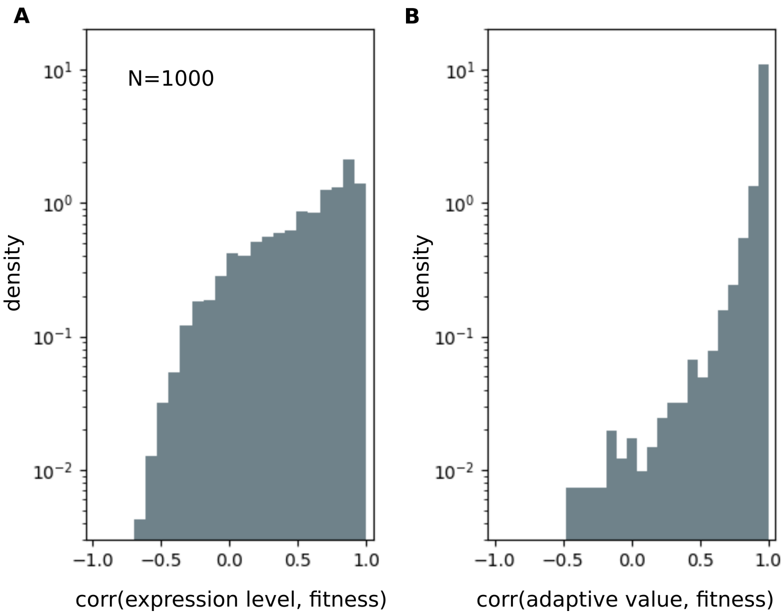


**FigS 19: Histograms for correlations between expression level, adaptive value and fitness trajectories for populations where the fitness threshold of 0.1 was crossed in the high mutation rate regime.** (A) correlations between expression level and fitness trajectories. (B) correlations between adaptive value and fitness trajectories.

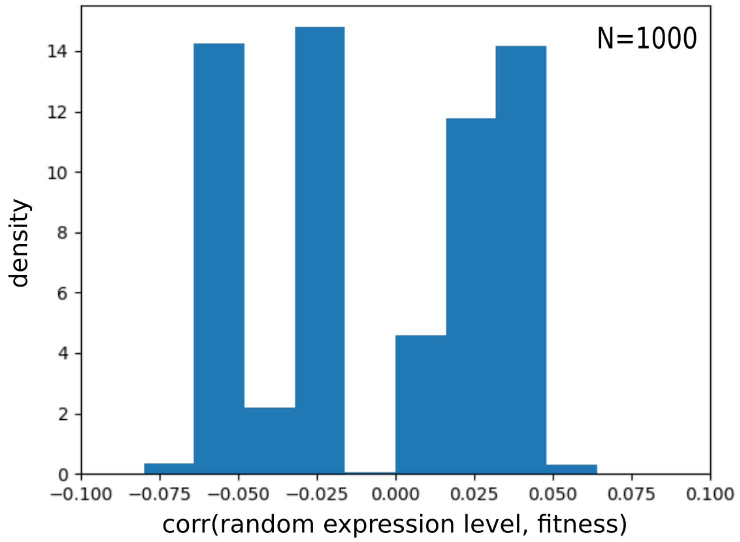


**FigS 20: Histogram for correlations between random expression level ($E^{rand}$) and fitness trajectories for populations where the fitness threshold of 0.1 was crossed in the high mutation rate regime.** We drew mutational effects on expression level ($\Delta E^{rand}$) from the same power law distribution used in the main simulations for N=1000 individuals for 1000 time steps. Here, no selection was involved in the update of expression levels, and for some individual i, $E^{rand}_{t+1}(i) = E^{rand}_t(i) + \Delta E^{rand}(i)$; we reset $E^{rand}(i)$ to 0.001 whenever $E^{rand}_{t+1}(i) < 0.001$. $E^{rand}(i)$ never crossed 1, which represents the highest possible expression level. While the trajectories of $E^{rand}$ and fitness show very little correlation, expression level trajectories obtained from the simulation are highly correlated (see FigS.21(A)).

Maarten Koornneef and Ben Scheres. Arabidopsis thaliana as an experimental organism. *e LS*, 2001.

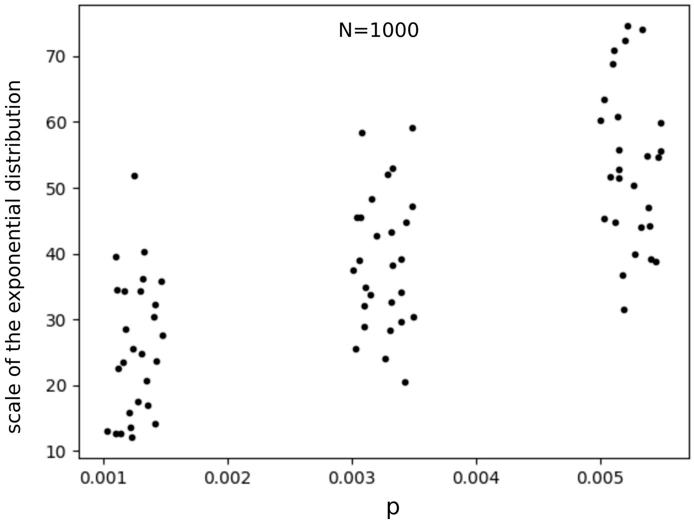


**FigS 21: Scale of $\Delta A$ exponential fits as a function of the model parameter $p$ in the high mutation rate regime.** Each point represents a parameter set; to obtain the fit, $\Delta A$ values were pooled from all time steps across all individuals across all replicates. The confidence intervals obtained for each fit through bootstrapping are too narrow to be visible in the figure.

Hongan Long, David J Winter, Allan Y-C Chang, Way Sung, Steven H Wu, Mariel Balboa, Ricardo BR Azevedo, Reed A Cartwright, Michael Lynch, and Rebecca A Zufall. Low base-substitution mutation rate in the germline genome of the ciliate tetrahymena thermophila. *Genome biology and evolution*, 8(12):3629–3639, 2016.

Yuan O Zhu, Mark L Siegal, David W Hall, and Dmitri A Petrov. Precise estimates of mutation rate and spectrum in yeast. *Proceedings of the National Academy of Sciences*, 111(22):E2310–E2318, 2014.

Heewook Lee, Ellen Popodi, Haixu Tang, and Patricia L Foster. Rate and molecular spectrum of spontaneous mutations in the bacterium escherichia coli as determined by whole-genome sequencing. *Proceedings of the National Academy of Sciences*, 109(41):E2774–E2783, 2012.

Ashley Farlow, Hongan Long, Stéphanie Arnoux, Way Sung, Thomas G Doak, Magnus Nordborg, and Michael Lynch. The spontaneous mutation rate in the fission yeast schizosaccharomyces pombe. *Genetics*, 201(2):737–744, 2015.

Rob W Ness, Andrew D Morgan, Nick Colegrave, and Peter D Keightley. Estimate of the spontaneous mutation rate in chlamydomonas reinhardtii. *Genetics*, 192(4):1447–1454, 2012.

Way Sung, Abraham E Tucker, Thomas G Doak, Eunjin Choi, W Kelley Thomas, and Michael Lynch. Extraordinary genome stability in the ciliate paramecium tetraurelia. *Proceedings of the National Academy of Sciences*, 109(47): 19339–19344, 2012.

Gerda Saxer, Paul Havlak, Sara A Fox, Michael A Quance, Sharu Gupta, Yuriy Fofanov, Joan E Strassmann, and David C Queller. Whole genome sequencing of mutation accumulation lines reveals a low mutation rate in the social amoeba dictyostelium discoideum. *Plos One*, 2012.

Peter A Lind and Dan I Andersson. Whole-genome mutational biases in bacteria. *Proceedings of the National Academy of Sciences*, 105(46):17878–17883, 2008.

Peter D Keightley, Urmi Trivedi, Marian Thomson, Fiona Oliver, Sujai Kumar, and Mark L Blaxter. Analysis of the genome sequences of three drosophila melanogaster spontaneous mutation accumulation lines. *Genome research*, 19 (7):1195–1201, 2009.

Stephan Ossowski, Korbinian Schneeberger, José Ignacio Lucas-Lledó, Norman Warthmann, Richard M Clark, Ruth G Shaw, Detlef Weigel, and Michael Lynch. The rate and molecular spectrum of spontaneous mutations in arabidopsis thaliana. *science*, 327(5961):92–94, 2010.

Katharina B Böndel, Susanne A Kraemer, Toby Samuels, Deirdre McClean, Josianne Lachapelle, Rob W Ness, Nick Colegrave, and Peter D Keightley. Inferring the distribution of fitness effects of spontaneous mutations in chlamydomonas reinhardtii. *PLoS biology*, 17(6):e3000192, 2019.

Eeshit Dhaval Vaishnav, Carl G de Boer, Jennifer Molinet, Moran Yassour, Lin Fan, Xian Adiconis, Dawn A Thompson, Joshua Z Levin, Francisco A Cubillos, and Aviv Regev. The evolution, evolvability and engineering of gene regulatory dna. *Nature*, 603(7901):455–463, 2022.

Jean-Pierre Eckmann and Tsvi Tlusty. Dimensional reduction in complex living systems: where, why, and how. *BioEssays*, 43(9):2100062, 2021.
